# Allopolyploidy enhances survival advantages for urban environments in the native plant genus *Commelina*

**DOI:** 10.1101/2024.06.24.600338

**Authors:** Hina Shimomai, Nakata Taichi, Koki R. Katsuhara, Seiji Kato, Atushi Ushimaru, Nobuko Ohmido

## Abstract

**Background and Aims:** Urbanization-induced environmental changes have profound effects on the geographical distribution limits in natural plant species. Polyploidization, an influential dynamic genome change, may determine survival potentials of plant species in urban environments. This study focused on the native plants, *Commelina communis* L. (Cc) and closely related subspecies, *C. communis* f. *ciliata* (Masam.) Murata (Ccfc) which have different chromosome numbers (e.g. Cc: 2n = 88, Ccfc: 2n = 46). The aim is to investigate polyploidization effects on natural plant distribution in urban environments.

**Methods:** The geographical distribution across urban-rural gradients was investigated at a total of 218 sites in Japan. Stomata size and density were measured and compared between Cc and Ccfc. Flow cytometry was used to determine genome size and polyploidy. Chromosome karyotyping was performed by using the GISH method.

**Key results:** Urban areas were exclusively dominated by Cc, while Cc and Ccfc coexisted in rural areas. Cc had larger and fewer stomata and more than twice genome size than Ccfc. GISH results indicated that Cc possesses Ccfc and another unknown genome, suggesting allopolyploidy.

**Conclusions:** These results show that the ploidy difference affects the geographical distribution, the stomata traits, and genome size between Cc and Ccfc. In addition, GISH results indicate that Cc has Ccfc and another unknown genome, suggesting Cc is an allopolyploid and these two species of the genus *Commelina* are distinct. Therefore, not only polyploid but also allopolyploid contributes to Cc to enhance survival potentials in urban environments compared to Ccfc. This is the first investigation to clarify the distribution difference related to urban environments, the difference in stomata traits and genome size, and to conduct chromosome composition in *Commelina* species.

## INTRODUCTION

One of the key questions in ecology is what factors limit the geographical distribution of plant species. Since the 20th century, anthropogenic pressures in natural and semi-natural environments have posed new challenges for plant species, of which urbanization is one of the main factors that alter abiotic and biotic environments of diverse habitats within and around cities (Pickett *et al*., 2001; Grimm *et al*., 2008; Johnson *et al*., 2015; Szulkin *et al*., 2020). In response to these new environments, some plants can survive and expand their geographical ranges, while others are limited in their expansion in urban areas. It is necessary to investigate the important factors that determine the geographical distribution limits of plant species in urban environments. Polyploidization, one of the most widespread and influential genomic dynamic phenomena, may lead to the potential difference in natural plants (Van Drunen and Johnson, 2022).

Polyploidization is an important for acclimatization to variable environments. Generally, polyploids, with more than two complete chromosome sets, have a higher tolerance to drought, rise in temperature, and salty conditions than diploids (Maherali *et al*., 2009; Chao *et al*., 2013; Godfree *et al*., 2017). Tossi *et al*. (2022) reviewed the tolerance to abiotic and biotic stresses as well as disease resistance can be enhanced by anatomical and physiological changes due to either natural or synthetic polyploidization, which may correlate positively with plant growth and net production. These tolerances of environmental stresses may benefit plants in urban areas (Johnson *et al*., 2015) and lead to expanding their habitats. At least, allopolyploids in *Veronica* subsection *Pentasepalae* differentiate their ecological distribution depending on climatic variables (Rojas-Andrés *et al*., 2019). However, there is almost no research that has verified the relationship between polyploidy and urban environments in naturally distributed species. An opinion paper summarizes several questions about the polyploidy–urban adaptation relationship (Van Drunen and Johnson, 2022). For instance, it is still unknown whether polyploids have a higher tolerance to stressful conditions and greater fitness in urban environments than closely related diploids, or not. It also remains unclear whether polyploids have larger populations in urban areas than in nonurban areas and whether this trend is consistent across different urban areas. Van Drunen and Johnson (2022) mentioned that urban environments may provide opportunities for neopolyploid establishment. However, already established polyploids in the system of native plant species have not been sufficiently discussed. Therefore, it is worth verifying these questions in several urban areas in wild polyploidy plants.

*Commelina communis* L. (Cc) and its forma, *C. communis*. f. *ciliata* (Masam.) Murata (Ccfc) are an appropriate model system to examine the relationship between polyploidization and urban environments using closely related subspecies including established polyploids. Their morphological and ecological traits entirely resemble each other: Cc and Ccfc have quite similar plant size, leaf size and morphology, flower size, shape, and flowering phenology (Katsuhara and Ushimaru, 2019; Fig. 1), also the habitat preference of Ccfc largely overlaps with that of Cc in rural areas (Katsuhara and Ushimaru, 2019). In urban areas, however, it is unknown how Cc and Ccfc are distributed. Furthermore, previous studies reported that Cc and Ccfc have great variations in chromosome number suggesting ploidy differences between them (Fukumoto, 1979; Fujishima, 1981, 2003, 2010).

**Fig. 1.**
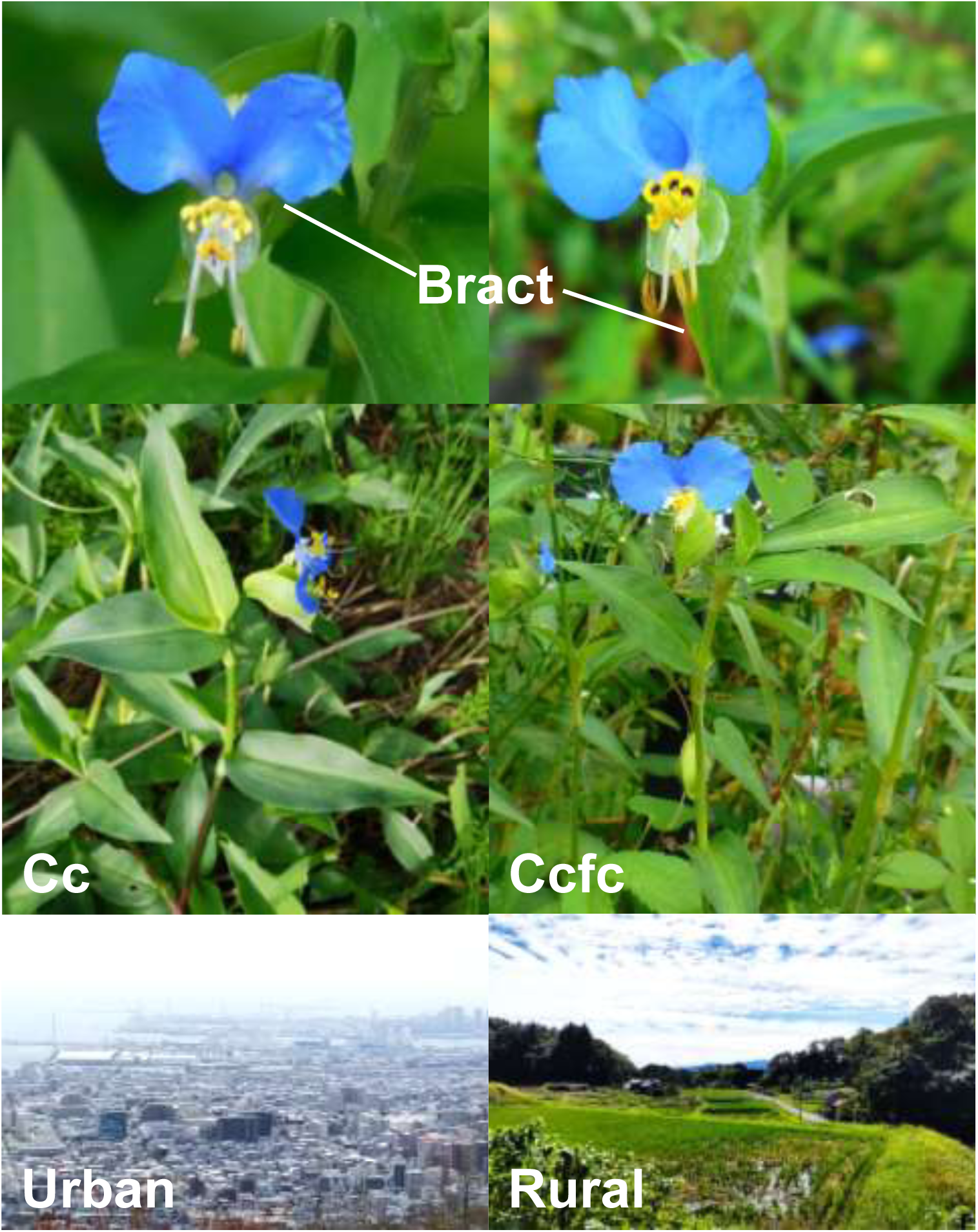
Flower (upper) and whole plants (middle) of *Commelina communis* (Cc) and *C. communis* f. *ciliata* (Ccfc) in urban and rural areas. Although Ccfc (right) is morphologically distinguished with Cc (left) due to bract hairs and shorter stamens (Katsuhara *et al.,* 2019), their morphological traits are highly similar. Urban area (lower, left) is occupied by developed lands such as concretes and buildings, whereas rural area (lower, right) is covered with natural fields such as rice fields and forests.

Although overall morphologies between Cc and Ccfc are highly similar, several studies showed that polyploidization causes increased sizes of cells or organs, such as flowers, shoots, leaves, pollens, and stomata (Van Laere *et al*., 2011; Sattler *et al*., 2016; Pei *et al*., 2019). Stomata are an important part of the leaf epidermis and its morphological traits can be sensitive to environmental changes (Zheng *et al*., 2015). Especially stomata size and stomata density are directly influenced by plant photosynthesis and transpiration (Flexas and Medrano, 2002; Flexas *et al*., 2002; Lawson and Blatt, 2014). At least, lower stomata density is reported to reduce the transpiration rate, leading to adaptation to drought conditions (summarized by Tossi *et al*., 2022). Furthermore, in urban high-temperature conditions, it has been reported that the stomata density increased, while the stomata area decreased significantly in tree leaves (Zhu *et al*., 2018). Therefore, understanding the relationship between polyploidization effects and urban adaptation through stomata traits in *Commelina* species provides an optimal approach to exploring the advantages of polyploidy plants in urban environments.

Polyploids are classified into two main categories by how they have arisen: simply homologous genome duplication (autopolyploids) or hybridization including two or more homoeologous genomes (allopolyploids). Allopolyploid populations are suggested to adapt to new conditions more quickly than closely related diploid populations, based on the comparison of climatic niches between allopolyploid vascular plants and their parental diploids (Baniaga *et al*., 2020).

Furthermore, white clover (*Trifolium repens*), a common allotetraploid forage crop, is reported to achieve niche expansions through allopolyploidization (Griffiths *et al*., 2019). To discuss plant distribution associated with polyploidy, it is important to clarify whether the target species are auto- or allopolyploid. Considering that Cc and Ccfc do not sexually cross with each other (Fujishima, 2010; Katsuhara and Ushimaru, 2019), it is difficult to verify the ploidy classification by chromosomal pairing. Cytogenetic approaches such as chromosome karyotyping and genome organization analysis will provide evidence of ploidy contracts.

In this study, we investigated the polyploidy–urban adaptation relationship using the genus *Commelina*, as a system of native plant species. For a better understanding of the urban distribution of natural plants relating to polyploidization, we tested the consistent geographical distribution patterns of Cc and Ccfc in urban–rural gradients of six urban areas including two world megacities, and analyzed the stomata, pollen, and genome sizes, chromosome numbers, and compositions, addressing the following questions: (1) Do the distribution patterns of Cc and Ccfc plants change along the urban–rural gradients and exhibit consistent trends across the cities? (2) Are ploidy traits different between Cc and Ccfc and do they relate to urbanization? (3) Is polyploid classification effective in understanding plant distribution in urban environments? We discuss the results considering the relationships between polyploidization and urban adaptation in the *Commelina* species.

## MATERIALS AND METHODS

### Study Commelina species

The genus *Commelina* is the largest genus of Commelinaceae with ca. 201-205 species (Lee *et al*., 2017; Royal Botanic Gardens, Kew: Plants of the World Online) and nine species of *Commelina* are known from eastern Asia to southeast Asia (Ridley, 1924; Hong and De-Filipps, 2000). According to the phylogenetic research of *Commelina* species, *Commelina* has two major clades; the first clade including *C. communis* (Cc), and the second clade of *Commelina benghalensis* L. (Cb) whereas its genome size and chromosome were used as an exotic species in our study (Lee *et al*., 2017). *Commelina communis* (Commelinaceae) is an annual, well-known ruderal plant widely-distributed in Asia (Fujishima, 2003), especially in eastern and northern Southeast Asia (Ohwi, 1975; Ushimaru *et al.,* 2014), and introduced in parts of Europe and North America and known as Asian dayflower (Royal Botanic Gardens, Kew: Plants of the World Online). It grows commonly in rice fields and on roadsides and riverbanks widely in Japan (Ohwi, 1975; Ushimaru *et al*., 2014). *Commelina communis* f*. ciliata* (Ccfc) is distinguished by *C. communis* (Cc). Ccfc has bract hairs and short stamens, while Cc has no bract hairs and long stamens (Katsuhara *et al.,* 2019; Fig. 1). Cc and Ccfc are mainly selfing plants and have a largely overlapped preference for their habitats (Katsuhara and Ushimaru, 2019). Furthermore, both Cc and Ccfc exhibit great variations in chromosome number; 2n = 86, 88, and 90 (Cc) and 2n = 44, 46, 48, 50, and 52 (Ccfc). The main chromosome number is 88 for Cc, 46 or 48 for Ccfc (Fukumoto, 1979; Fujishima, 1981, 2003, 2010). In addition, Cb in Japan is 2n = 22 (Fujishima, 2007).

### Study sites

A total of 218 sites (41, 30, 42, 37, 40, and 28 from Shimane, Okayama, Hyogo, Osaka, Kyoto, and Tokyo prefectures, respectively) were examined in 2021 and 2022 (Supplementary data Table S1). The Tokyo and Kyoto–Osaka–Kobe metropolitan areas are the world’s largest megacities with approximately 39 and 19 million residents, respectively (United Nations, 2019). In 2021, 27 Cc plants and 16 Ccfc plants were collected, while 24 Cc plants and 14 Ccfc plants were sampled in 2022 (Supplementary data Table S2). In previous paper of Katsuhara *et al*. (2019), a single voucher specimen of Cc, Ccfc, and Cb collected from Hyogo were submitted in the Osaka Museum of Natural History in Japan; accession number: OSA300034 (Cc), OSA300037 (Ccfc), and OSA300036 (Cb). In this study, we identified plant samples based on these specimens. As they were collected from roadsides and rice fields, no permits were required at most collection sites, while official permissions were obtained at three parks in Tokyo: Tokyo Metropolitan Nogawa Park, Senzokuike Pond Park, and Saneatsu Park (Tokyo, Japan). After plant collections, 51 Cc and 30 Ccfc plants were transplanted separately into pots (10 cm diameter × 12 cm height; approx. 1.2 L) in the common garden of Kobe University (34°73′N, 135°23′E).

### Categories of the geographical distribution

The geographical distribution of Cc and Ccfc populations was examined to assess the effects of urbanization on these *Commelina* species (Supplementary data Table S1). For each site, we classified the ratio of the *Commelina* species distribution as follows; 1: 0% of Cc, 2: 1-25%, 3: 25-50%, 4: 50-75%, and 5: 100%, depending on their relative abundances. 0% and 100% show that no Cc and no Ccfc populations was found at each site, respectively. These data were also summarized into three levels: Ccfc only (1), Co-existence of Cc and Ccfc (2, 3, 4), and Cc only (5).

### Categories of land use

The cumulative areas of developed land within radii of 500 m and 1000 m from the centre of each study area were quantified using the QGIS (Ver. 2.0, QGIS Development Team). Land use and land cover data were obtained from a high-resolution map (2018–2020) produced by Japan Aerospace Exploration Agency (JAXA, 2022, 10 m equivalent resolution version 21.11). The data were divided into four categories: water, agricultural land, forest, and developed land.

### Stomata trait measurements

In general, higher ploidy plants have larger stomata size and lower density compared to lower ploidy plants, which is supported by the study of *Arabidopsis* autotetraploids and its diploids as model plants (del Pozo and Ramirez-Parra, 2014). We investigated whether the differences related to polyploidy such as stomata size and density are exhibited between Cc and Ccfc, 36 plants for 36 sites (one plant per site) were selected and one leaf per plant was collected (Supplementary data Table S3): in 2022, from three regions (Kyoto, Hyogo, and Shimane), fresh leaves of 6 plants of Cc and 3 plants of Ccfc in urban and rural areas were collected directly and stored at 4L for the analysis of stomata traits and fresh leaf samples of Tokyo were collected and used after transplanting the plant samples. The leaf samples were collected from the upper of each plant and every base of the leaf was separated for stomata trait measurements. The stomata images from the abaxial epidermis layers from the leaves were observed under a light microscope BX60 (Olympus, Tokyo, Japan) followed by taking photos imaging by CCD camera (SPOT-RT3, Meyer Instruments, Inc.) and recording the length of 1080 stomata and density per 0.26 mm^2^ using ImageJ software (Rasband *et al*., 1997-2018).

### Flow cytometry analysis

The fresh leaves from Kyoto and Shimane were kept at 4L and used for the analysis of flow cytometry in 2021 (Supplementary data Table S4). Flow cytometry was performed to determine genome size and ploidy between two related species, following the previous protocol (Beckman Coulter, application note 6) with some modifications. Briefly, the fresh leaf samples stored at 4L for a week were immersed in 1.2 mL chopping buffer (1.0% Triton X-100, 140 mM 2-Mercaptoethanol, 50 mM NaHSO_3_, 50 mM Tris hydrochloric acid, and 25 µg/mL Propidium iodide) in a 6 cm petri dish and chopped with a new sharp razor blade. After enough chopping of the suspension in the buffer, the nuclei suspension was passed through a 20 µm nylon mesh filter, then placed on ice with more 0.6 mL of chopping buffer. The nuclei suspension was centrifuged at 1,000 rpm for 2 min, followed by removingthe supernatant. After more 0.4 mL of chopping buffer was added and filtered through a filter into the test tube, the samples were analyzed using a flow cytometer (BD FACSAria TM L Cell Sorter). The genome size control was a leaf sample of *Oryza sativa*, subsp. japonica cv Nipponbare, treated in the same way. The genome size values were converted from the original data and compared to the genome size of *O. sativa* as a control (389 Mb/1C, International Rice Genome Sequencing Project, 2005).

### Chromosome preparation

Chromosome specimens of Cc, Ccfc, and Cb were prepared for chromosome number analysis, using a previous protocol (Ohmido *et al*., 2013) with some modifications. Root tips for chromosome number and karyotype analysis were collected from plants in each area grown in the garden in 2022. Firstly, root tips were excised and pretreated in a mixture of 2 mM 8-hydroxyquinoline and 0.025% colchicine (4:1) at 18-21L for 3-4 h. After pretreatment, the samples were washed in distilled water, followed by fixing in Carnoy’s solution (100% (v/v) ethanol: 100% (v/v) acetic acid = 3:1) at - 20L before use for future experiments. After washing with distilled water, they were digested in an enzyme mixture of 1% Cellulase Onozuka RS (Yakuruto Ltd, Tokyo, Japan), and 2.5% Pectolyase Y-23 (Seishin Ltd, Tokyo, Japan). The roots were vacuumed for 5 min in a desiccator at -0.1mPA and incubated for 40-70 min at 37L. The root tips were added 10-30 µL of Carnoy’s solution (Ethanol: acetic acid = 3:1) and squashed speedily with forceps. The slides on which chromosome samples are observed under a microscope then were stored at 4L to use for further GISH.

### Genomic in situ hybridization (GISH)

For analysis of auto- or allopolyploid, chromosome karyotyping with genomic fluorescence *in situ* hybridization (GISH) is the practical method for chromosomal mapping; total genomic DNA is used for GISH to know the ancestral genomes (Schwarzacher *et al*., 1989, Ohmido *et al*., 2013; Heslop-Harrison *et al*., 2023). Genomic DNA was isolated from fresh leaves using the DNeasy Plant Mini Kit (QIAGEN, Hilden, Germany). Ccfc genomic DNA was labeled indirectly by Nick translation mix (Roche, Versel, Swissland), with biotin-dUTP (Roche, Versel, Swissland). The GISH method using the genomic DNA of Ccfc was performed using well-prepared chromosome spreads according to the method described by Ohmido *et al*. (2010, 2013) with slightly modified. Chromosome spreads are treated by 100 ng/µL RNase treatment in 2×SSC at 37L for 30 min. After washing with 2×SSC, slides were pretreated in 1% Formaldehyde/PBS at RT for 4 min. After washing twice in PBS for 5 min each and dry-up. The hybridization mixture (50% Formamide, 1×SSC, 1% SDS, 10% Dextransulfate) containing 100-200 ng biotin labeled Ccfc genomic probe was denatured at 80°C for 10 min and kept on ice. Slides were denatured in hybridization buffer with the biotin-labeled genomic probe at 70L for 4 min and gradually decreased to 55L for 4 min. After overnight hybridization at 37L in a dark humid box and the slides were washed in the slides were washed in 20% Formamide/2×SSC at 42°C, three times for 5min each and 0.1×SSC at 42°C, three times for 5min each. Slides were incubated with Streptavidin Cy3 (Jackson Immuno. Res. Lab, USA) in 4×SSC at 37°C for 60 min. The chromosome samples were washed in 2×SSC at room temperature three times for 5 min each. After washing and air-drying, the chromosome samples were stained with 1 µg/ml 4,6-diamidino-2 phenylindole (DAPI) in Vectashield (Vector Laboratories). The slides were observed under a fluorescence microscope (BX60; Olympus, Tokyo, Japan) equipped with SPOT RT3 CCD camera. The GISH images were captured using blue, and green filter using standard Olympus filter sets. Image analyses (160 images for Ccfc, 154 for Cc, and 81 for Cb) were completed by ImageJ.

### Statistical analysis

All statistical analyses were performed using R version 4.2.1 (the R Development Core Team, 2022) with the “glmmTMB” package (Blooks *et al*., 2017) and/or multcomp” package (Hothorn *et al*., 2008).

Firstly, we examined the effects of urbanization on the presence/absence of Cc and Ccfc in each study site using the generalized linear mixed models (GLMM) with a binomial distribution and logit link functions. The presence of Cc/Ccfc for each site was incorporated as the response variable, the proportion of developed land area around the site and region identity (a category of the region: Tokyo, Kyoto, Osaka, Hyogo, Okayama, and Shimane) were explanatory variables, plant species identity and the proportion of developed land area were interaction terms, and sampling year (2021 or 2022) was random term. The lowest (AIC) model with the landscape variable of either spatial scale (500 m or 1000 m) was selected as the best model.

To examine the effect of the urban environments on the stomata size and density of Cc, the GLMM with a Gaussian distribution and identity link functions were performed. Mean stomata size per leaf was the response variable, whereas the ratio of developed land area around each site and study region identity were the explanatory variables, and study site identity was the random term. The differences in ploidy and altered stomata size and density between Cc and Ccfc were also examined by using GLMMs with a Gaussian distribution and identity link functions. Stomata size was the response variable, and plant species identity (Cc/Ccfc) with study region identity and leaf area were the explanatory variables whereas study site identity was the random term. Stomata density was analyzed in the same way as stomata size. Leaf area and collection date were analyzed with stomata size to consider the effect. Multiple comparison was performed for plant species identity and the region identity by a Tukey method. Each model with the lowest AIC was selected for these analyses.

## RESULTS

### Geographical distribution differences in Cc and Ccfc

The geographical distribution of *C. communis* (Cc) and closely related subspecies *C. communis.* f*. ciliata* (Ccfc) were investigated to clarify the distribution pattern in urban-rural gradients. In rural areas where forests and agricultural fields were largely maintained, both Cc and Ccfc were sympatrically distributed although some sites such as sites in Shimane had only Cc (Fig. 2D). On the other hand, in urban areas with larger developed lands, Cc was exclusively distributed and almost no Ccfc was found. Only Cc sites were observed widely in urban and rural areas, while co-existence and only Ccfc sites were found only in rural areas. It should be noted that a few co-existence sites were found even in urban areas (Fig. 2A, 2B, and 2C). Furthermore, the relationship between the presence of Cc or Ccfc and the developed lands was shown in Fig. 3, and Supplementary data Table S5. In all cities, the presence of Ccfc decreased significantly with increasing the proportion of surrounding developed land areas, whereas Cc was consistently present across urban-rural gradients. Thus, in all the cities, the relative dominance of Cc was more frequent as the developed lands increased, while Ccfc disappeared (Supplementary data Fig. S1; Table S6).

**Fig. 2.**
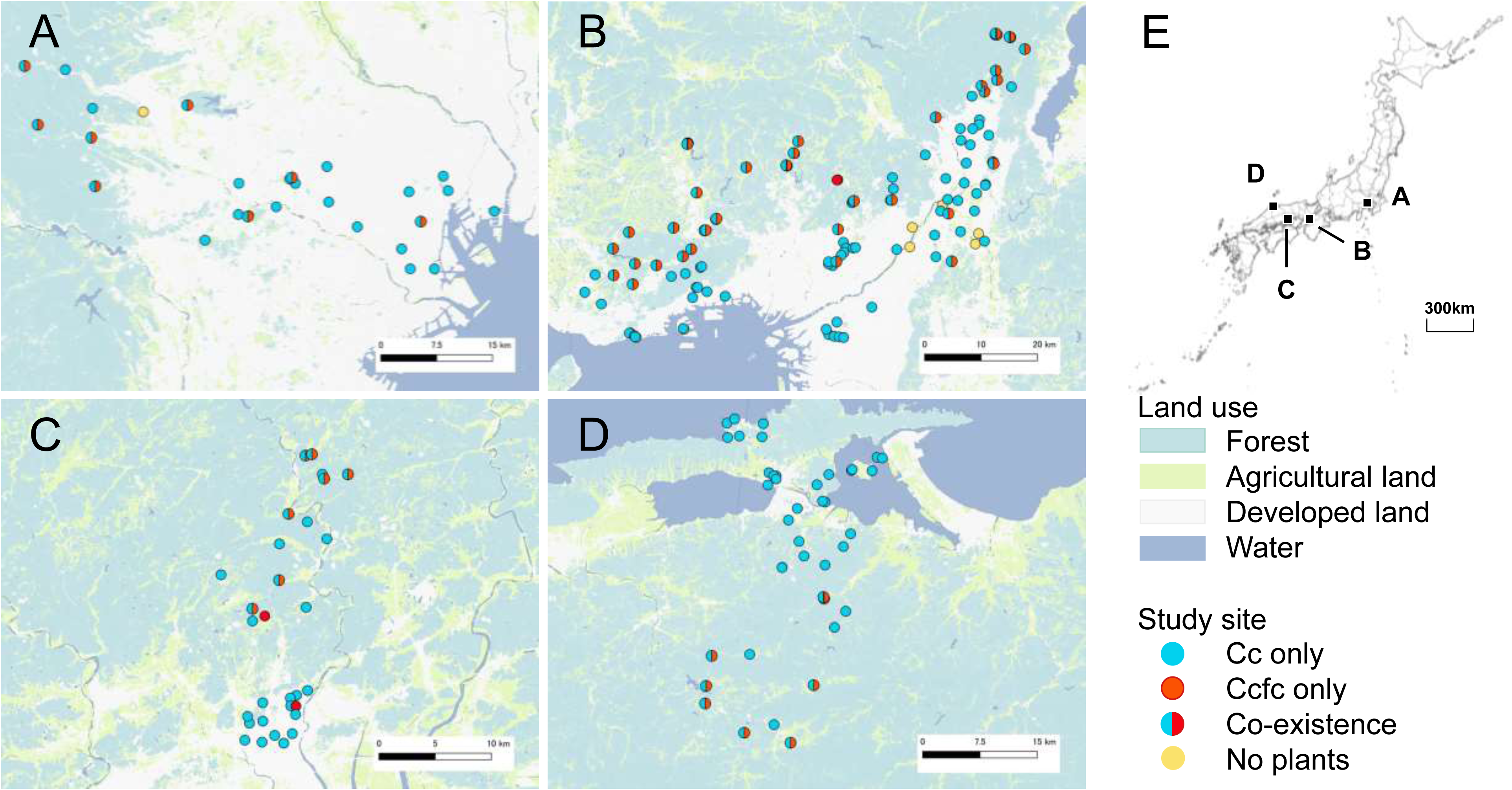
Geographic distribution of *Commemina communis* (Cc) and *C. c.* f. *ciliata* (Ccfc). (A) Study sites in Tokyo. (B) Study sites in Hyogo, Osaka, and Kyoto. (C) Study sites in Okayama. (D) Study sites in Shimane. (E) Overall study regions.

**Fig. 3.**
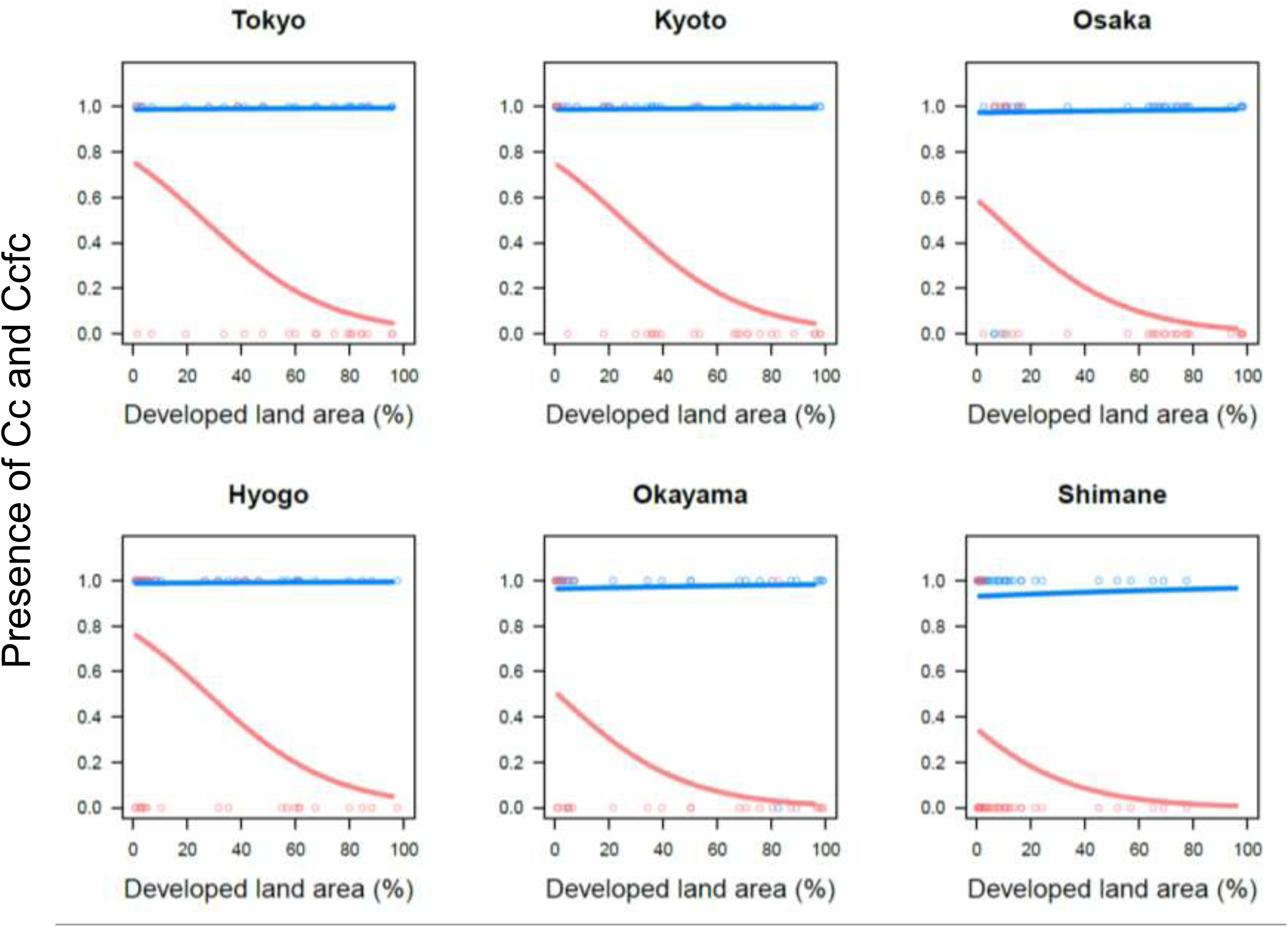
The relationships between Cc and Ccfc presence and developed land area. The horizontal axis indicates the percentage of developed land areas (%) in each site. The vertical axis indicates the presence of Cc and Ccfc in each developed land area. The regression lines of Cc (blue) and Ccfc (red) are illustrated by using GLMM dataset (Table S5). All the regression lines of Ccfc (red) are illustrated by the significant explanatory variable. Each plot means the presence/absence (1/0) of Cc (blue) and Ccfc (red).

### Stomata trait variations of Cc and Ccfc

Stomata size and density of Cc and Ccfc were measured and analyzed to clarify the effect of polyploidization by using the data collected from rural areas. The variation in stomata size and density along urban–rural gradients was also analyzed to understand the effect of urban environments on these variables by using only Cc data. Overall, the stomata length of Cc was significantly longer than that of Ccfc in all regions (*e.g*.The average of Cc stomata size was approx. 1.2 times longer than that of Ccfc one: 51.91±5.61 μm in Cc, 43.23±5.07 μm in Ccfc, Fig. 4; Supplementary data Table S7; Fig. S2). The stomata length of Cc varied among regions such as the existence of significant differences between Tokyo and Shimane, while that of Ccfc showed no significant differences within Ccfc individuals (Fig. 4; Supplementary data Table S8, S9). Stomata density of Cc was significantly lower than that of Ccfc (Supplementary data Table S7), however, the density varied among regions and within the same species due to the widely varied density of Ccfc in Kyoto and Cc in Tokyo (Supplementary data Table S8, S10). There was no significant relationship between stomata traits (size and density) and the developed land areas in Cc (Supplementary data Table S11; Fig. S3). From the analysis of the relationship between stomata traits, leaf area, or collection date (Supplementary data Table S11, S12), some significant differences in those factors were found such as in Tokyo regions (*e.g.* stomata size in Shimane and Hyogo compared to Tokyo). In addition, the pollen size of Cc was significantly larger than that of Ccfc (Supplementary data Table S13).

**Fig. 4.**
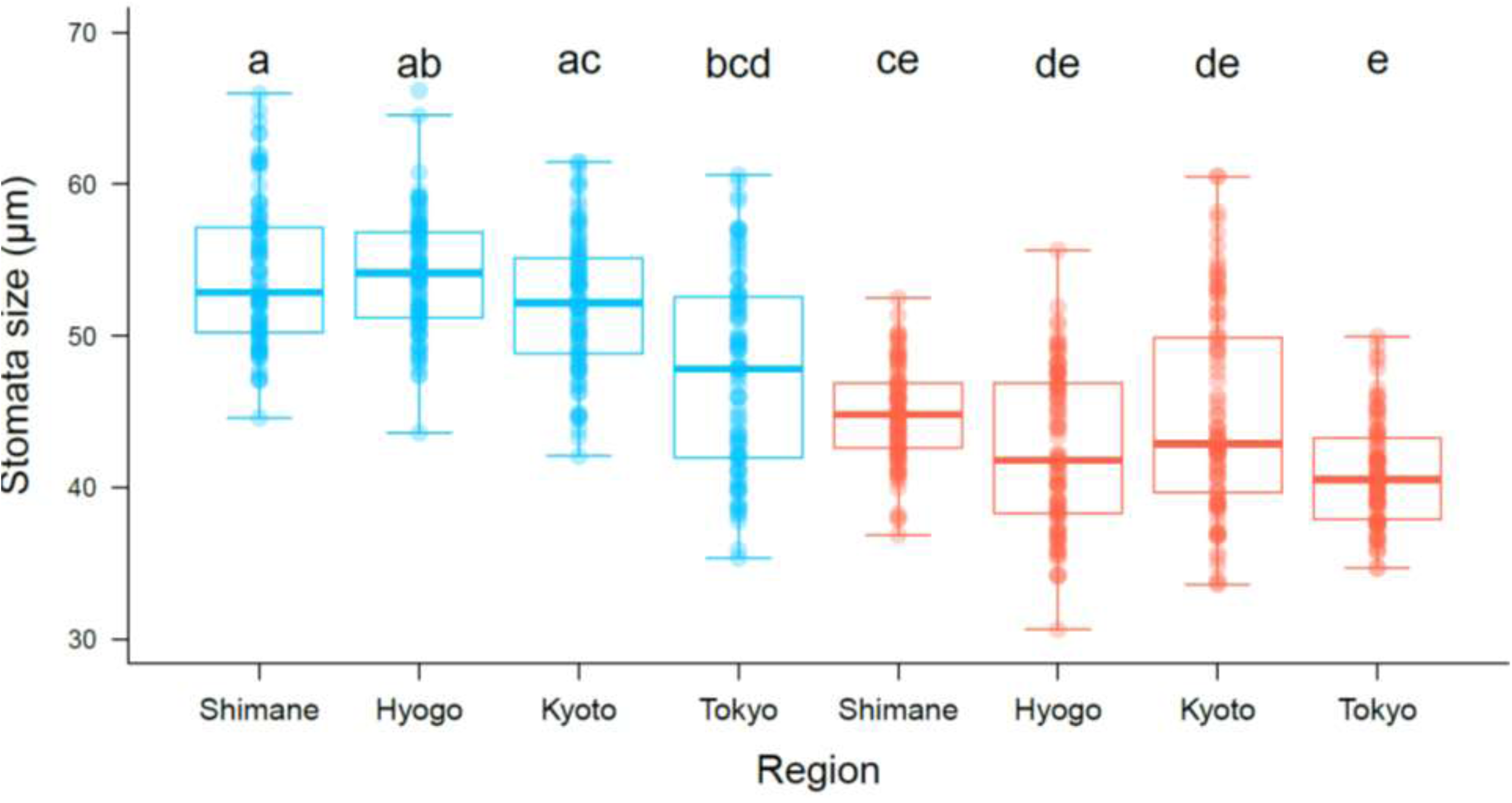
The comparison of Cc and Ccfc stomata size in regions. Blue and red boxplots show *Commelina communis* (Cc) and *C. c.* f*. ciliata* (Ccfc), respectively. Each plot means the raw data of stomata size. Different letters indicate the significant difference in species and regions (p<0.01).

### Genome content of Cc and Ccfc

The determination of genome size and ploidy to confirm the ploidy difference between Cc and Ccfc was performed by flow cytometry. As shown in Table 1, while the DNA content of Ccfc displayed 2973.62 Mb/1C, the Cc DNA content was 6969.83 Mb/1C (both data were averages of three samples in Kyoto). The average genome size of Cc was more than twice that of Ccfc both in Shimane and Kyoto. However, there was no significant difference between Shimane and Kyoto on the same taxa, Cc and Ccfc, respectively. These data suggested that Cc is polyploidy compared to Ccfc and both genome size of Cc and Ccfc do not change depending on geographical regions. However, the ploidy levels of Cc and Ccfc cannot be concluded from these data due to intraspecific chromosome variation (Fukumoto, 1979; Fujishima, 1981, 2003, 2010) and the diversity in the basic chromosome number as discussed below. Also, the DNA content of *C. benghalensis* (Cb) was 2312.63 Mb/1C, which was not half of Ccfc despite its chromosome number.

**Table 1.** Genome size of Cc and Ccfc measured by flow cytometry. *Oryza sativa* was used as a control (389Mb/1C, 2n = 2x = 24). *C. benghalensis* is one of the exotic species in Japan. Only one plant was measured genome size in *C.benghlensis* and *O. sativa*, respectively.

| Species | Region | Average (Mb/1C) | Average (pg/1C) | Chromosome number |
| --- | --- | --- | --- | --- |
| *C. communis* | Shimane | 6820.35±450.88 | 6.97±0.46 | 88 |
| *C. c. f. ciliata* | Shimane | 2747.93± 39.04 | 2.81±0.04 | 46 |
| *C. communis* | Kyoto | 6969.83±750.73 | 7.13±0.77 | 88 |
| *C. c. f. ciliata* | Kyoto | 2973.62±348.13 | 3.04±0.36 | 46 |
| *C. benghalensis* | Kyoto | 2312.63 | 2.36 | 22 |
| *Oryza sativa* (control) | — | 389 | 0.40 | 24 |

### Chromosome numbers and genome composition by genomic in situ hybridization (GISH)

A comparison of chromosome number and composition of Cc, Ccfc, and Cb is shown in Fig. 5. Cc, Ccfc, and Cb showed 88, 46, and 22, respectively. Focusing on the chromosome size, Ccfc and Cb were larger than Cc despite condensed phases (Supplementary data Fig. S4). In addition, several morphological differences in chromosomes were observed among the three species. Cc chromosomes had more chromosome knobs at the end and in the middle, and elongated shapes, compared to Ccfc and Cb (Fig. 6A, 6D, 6G).

**Fig. 5.**
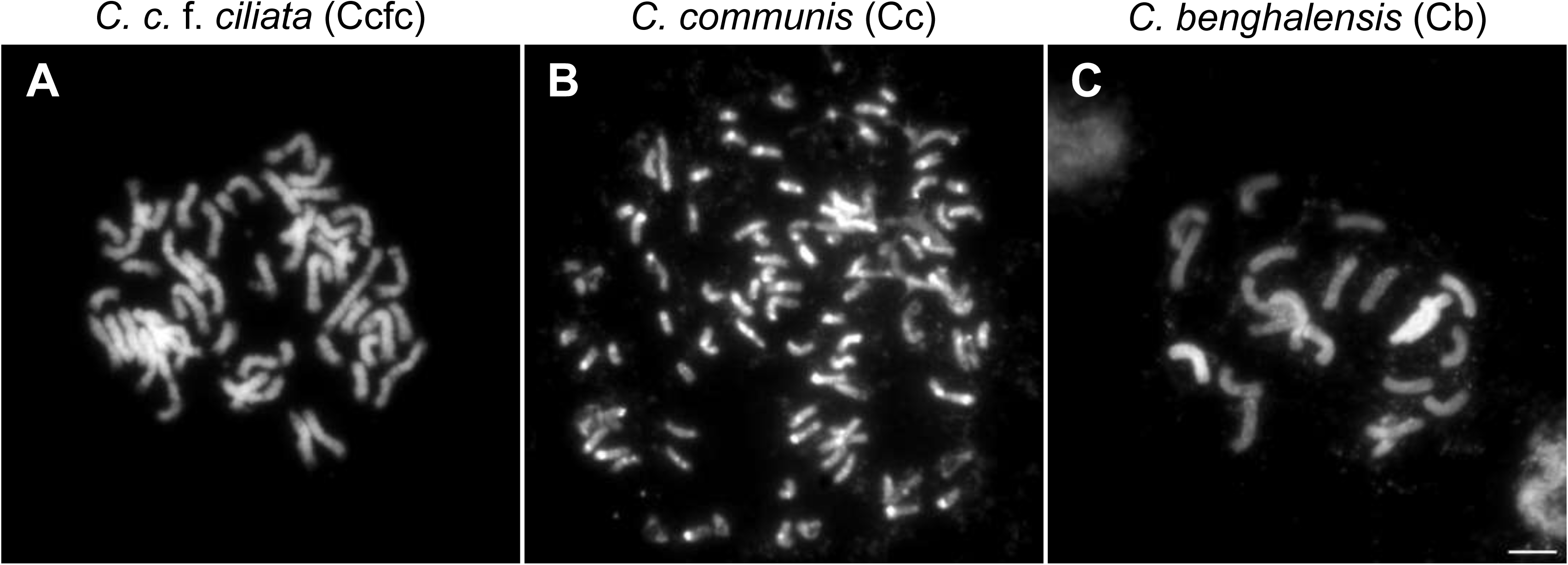
Mitotic metaphases of Cc, Ccfc, and Cb. (A) Ccfc chromosomes stained by DAPI (2n = 46). (B) Cc chromosomes stained by DAPI (2n = 88). (C) Cb chromosomes stained by DAPI (2n = 22). Scale bar: 5 µm.

**Fig. 6.**
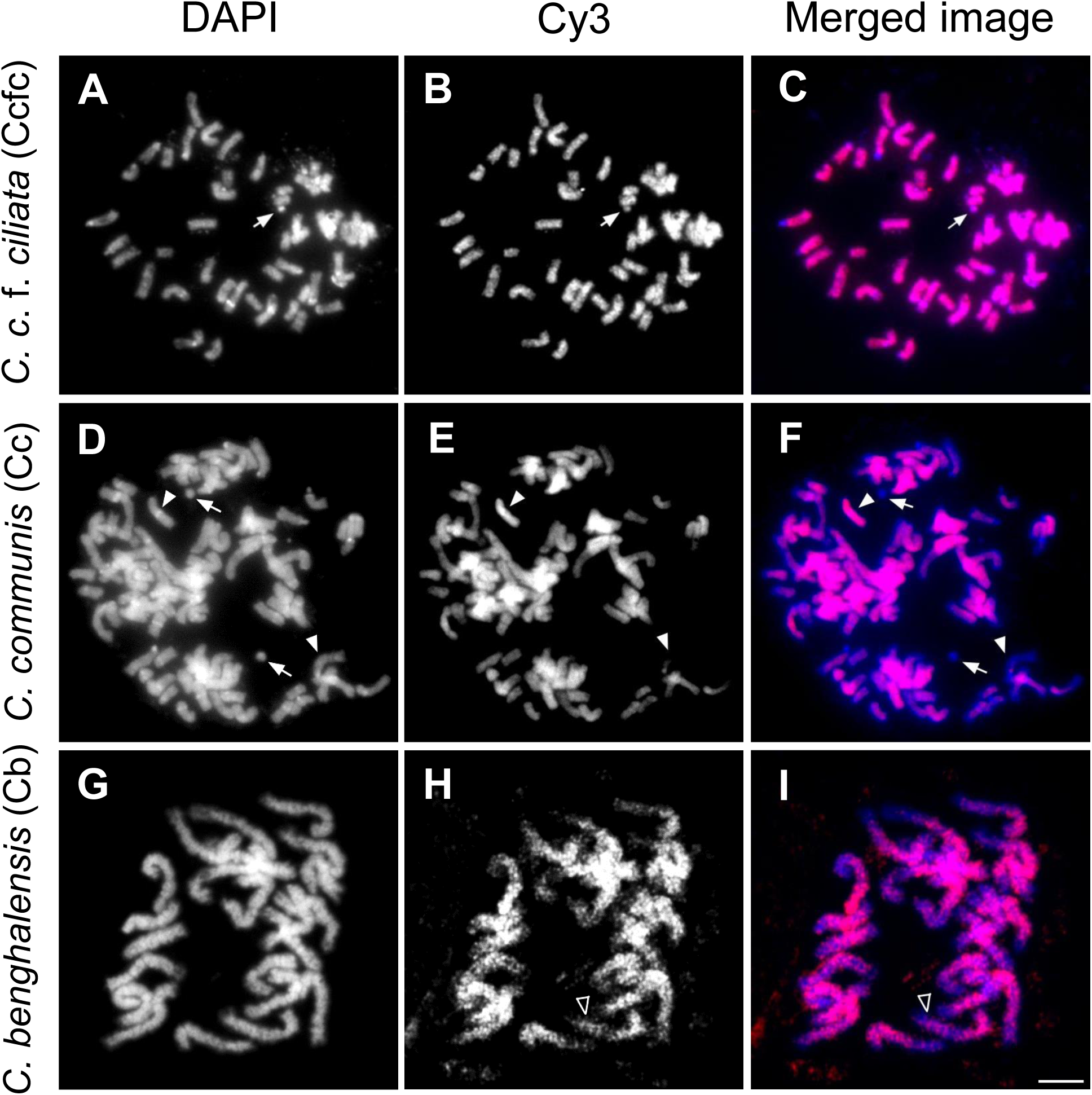
Mitotic metaphases of Cc, Ccfc, and Cb after GISH using genomic Ccfc DNA. (A) Ccfc chromosomes stained by DAPI. (B) Ccfc DNA signals labeled by Cy3. (C) Ccfc DNA signals (red) merged on Ccfc chromosomes. (D) Cc chromosomes stained by DAPI. (E) Ccfc DNA signals labeled by Cy3. (F) Ccfc DNA signals (red) merged on Cc chromosomes (blue). Ccfc genomic DNAs localized on a half part of Cc chromosomes. Nucleolus organizer region (NOR) does not appear Ccfc signals. (G) Cb chromosomes stained by DAPI. (H) Ccfc DNA signals labeled by Cy3. (I) Ccfc DNA signals (red) merged on Cb chromosomes (blue). Arrows demonstrate NOR. Arrowheads (white frame) indicate band pattern of the signals. Arrowheads (white) indicate the signal patterns are different depending on each chromosome. Scale bar: 5 μm.

Chromosome composition was performed using genomic DNA as genome marks to distinguish the genome components among Cc, Ccfc, and Cb at the chromosomal level (Fig. 6). Two species (Ccfc and Cb) were used for generating GISH probes. Firstly, Ccfc DNA probe showed strong and uniform signals on all Ccfc chromosomes (Fig. 6C). In contrast, the combination of the Ccfc DNA probe and Cc chromosome showed a quite different pattern: the Ccfc DNA signals appeared unevenly and showed both strong and weak fluorescent signals, although the Ccfc DNA signals were detected as well as on overall Cc chromosomes (Fig. 6F: Arrows and arrowheads). Furthermore, Ccfc DNA probe exhibited band signals on Cb chromosomes (Fig. 6I). Cb genomic DNA probes appeared as dot signals on several Ccfc chromosomes (Supplementary data Fig. S5C). When the Cb DNA was hybridized on Cc chromosomes, the signal appeared weakly on all chromosomes (Supplementary data Fig. S5F). This result suggests that Cc has Ccfc chromosomes and another set of chromosomes and that Cc is not an autopolyploid of Ccfc but an allopolyploid. As for Cb and the two species, the result indicates the difference in signal patterns.

## DISCUSSION

### Polyploid Commelina species dominates across urban and rural areas

These results showed that in urban areas *i.e.* stressful environments, Cc was exclusively dominant compared to Ccfc in every urban area. Only Cc can survive even in urban environments, while Ccfc prefers rural areas where the effect of urbanization is relatively low. Although Cc and Ccfc morphology traits are quite similar, Cc were consistently and exclusively dominant in all the study urban areas. As shown by flow cytometry, stomata traits, and chromosome numbers, Cc represents polyploidy characteristics. Polyploids usually have a broader distribution than their diploids (Liu *et al*., 2019). These results can answer the following questions: do polyploids have a larger population in urban environments than in nonurban environments? Is the trend consistent across cities? Polyploid Cc exhibits wider distributions and larger populations in all urban areas than closely related Ccfc. Concerning geographical distribution, high levels of ploidy are considered to enable plants to have a high invasive capacity leading to expanded distribution (Moura *et al*., 2021). Thus, our findings suggest that the polyploidization of *Commelina* species leads to wide distributions in urban areas including world megacities. However, it should be noted that Ccfc was found in some urban areas, which contradicts the overall results. Focusing on those sites in Tokyo, Ccfc appeared at large and semi-natural parks with high plant species richness (Fig. 2A). Those parks with Ccfc may have an important role as a refuge for original plants and insects (McIntyre, 2001; Goddard *et al*., 2010; Dohzono and Ushimaru, 2014). Although Ccfc can survive in such well-conserved urban areas, their populations might gradually decrease as the urbanized developed area increases. Allopolyploid enhances the potential for survival in urban environments.

This is the first investigation using the GISH method in *Commelina* species and the results indicate allopolyploid Cc (2n = 88) is not duplicated simply from Ccfc (2n = 46) but has a different genome composition (Fig. 6). It is also supported by the fact that Cc and Ccfc taxa do not sexually cross with each other (Fujishima, 2010; Katsuhara and Ushimaru, 2019) and different chromosome numbers in the genus *Commelina*. Although there are no reports showing auto- or allopolyploid of Cc, however, Katsuhara *et al*. (2019) demonstrated a high degree of genetic differentiation between Cc and Ccfc by principal component analysis (PCA), suggesting that Cc is not simply a duplicate of Ccfc. Analysis of microsatellite markers by Katsuhara *et al*. (2019) shows that 13 of the 15 markers overlapped in the range of amplicon sizes in cross-amplification testing between Cc and Ccfc, suggesting that they share a common chromosome. Katsuhara *et al*. (2019: Table S1) shows the results of testing Cc-SSR markers developed by Li *et al*. (2015). Only 7 out of 12 markers were amplified with the Ccfc genome. According to these reports and our results, Cc includes the Ccfc homoeologous chromosomes and is an allopolyploid. Different genome existence means having a larger genetic pool and enhances the genetic diversity of Cc than that of Ccfc, leading to such a high potential for environmental adaptation in Cc. Although there might be a possibility that one or more sets of additional chromosomes could carry functional multiple genes transcribing in the polyploid and have a potential role by expressing, not genome duplication (multiple genes), but an introgressed unknown genome could be associated with this added fitness. Previous studies showed that polyploidy was an essential key to the survival and establishment of the plant lineage during and after climatic upheavals (Estep *et al*., 2014; Cai *et al*., 2019). Since polyploidy is often associated with several potentials for survival such as high tolerance to stressful environments (Van de Peer *et al*., 2020), allopolyploidization may enable polyploids to have more stress tolerances in urban environments. In contrast, a recent study demonstrated that there was no evidence that allopolyploids have larger and more extreme geographical distributions than their ancestors, rather the distribution of allopolyploids highly overlapped with their ancestors (Mata *et al*., 2023). However, their study focused on climatic categories such as maximum or minimum precipitation and temperature. Our study proposes the new insight that allopolyploidy may provide advantages to expand a plant species on more microecological scales such as urban–rural gradients. Furthermore, polyploidy has also been reported to be associated with evolutionary dead end, low genetic diversity, and low speciation rates compared to the diploid ancestors or closely related subspecies (Mayrose *et al*., 2011; 2015). Alsayied *et al*. (2015) highlighted that the allotriploid species *Crocus sativus* (commonly known as saffron), grown worldwide, exhibits very limited genetic diversity. Through asexual reproduction (clonal propagation), the species leads to minimal genetic variation among cultivated saffron plants. The limited genetic diversity has implications for plant adaptability and disease resistance, presenting significant challenges for cultivation and conservation efforts. Cc has a higher prior selfing ability (Katsuhara and Ushimaru, 2019). Urban areas generally have fewer pollinators and increased selfing rates than rural areas (Ushimaru *et al*., 2014). The genetic diversity was not significantly different between urban and rural areas (Taichi *et al*., 2024). Cc in the same megacity showed little effect of urbanization on genetic structure. These indicate that polyploidy might reduce the negative impacts in urban environments on genetic diversity (Van Drunen and Johnson, 2022; Caizergues *et al*., 2024; Taichi *et al*., 2024). Urbanization had little effect on the genetic structure of the allopolyploid *Trifolium repens*. Allopolyploid with a larger genetic pool can also support maintaining genetic diversity in urban populations. Furthermore, Cc and Ccfc also have outcrossing traits producing single staminate and perfect flowers within a single bract. Many pollinators visit both flowers of Cc and Ccfc in the natural field (Ushimaru *et al*., 2009). According to these reports and our results, polyploidy enables Cc to maintain genetic diversity and the possibility of outcrossing leads to enhancing the diversity. Therefore, in the case of Cc, polyploidy may not be associated with an evolutionary dead end, but rather contribute to the expansion of this species as expanding on a global scale. In rural areas, co-existence may be associated with Cc having a Ccfc genome. This co-existence is also supported by the fact that Cc is less susceptible to reproductive interference from Ccfc due to its higher prior selfing ability than Ccfc (Katsuhara and Ushimaru, 2019).

Our finding that Cc populations dominate in urban areas is also consistent with previous ecological investigations since the wider distribution of Cc compared to Ccfc has been reported at least by Fujishima (2010), Ushimaru *et al*. (2014) and Katsuhara and Ushimaru (2019). Cc populations are widespread throughout the world (Royal Botanic Gardens, Kew: Plants of the World Online), although Ccfc is not found.

Survival advantages are regulated by factors including pollinators, flowering traits, and seed conditions such as the number of viable seeds produced and successful seed dispersal. In the comparison between Cc and Ccfc, there are no significant differences in seed production, pollinators, or flowering time, and their flower sizes are highly overlapped (Katsuhara and Ushimaru, 2019). The number of flowers was not significantly different between Cc and Ccfc under the same conditions (Murakami, unpublished data). Although Ccfc can be distinguished by bract hairs and the structure angle of stamen, these morphological characteristics are unlikely to be survival disadvantages. The biomass of Cc was significantly greater than that of Ccfc under the controlled conditions (Murakami, unpublished data), however, it is difficult to compare the biomass in the natural fields as the biomass is influenced by many other factors. Therefore, stomata or other metabolic traits related to ploidy levels observed in this study may lead to survival advantages for Cc compared to Ccfc. To compare the survival ability between Cc and Ccfc, future studies should examine the germination and recovery rates of Cc and Ccfc.

The distribution results suggested that polyploids distribute larger and have a higher tolerance to urban environmental stresses. In urban Cc habitats, compared to rural habitats, there are higher levels of soil pH, drought, shaded, and pollinator-limited conditions (Ushimaru *et al*., 2014; Taichi and Ushimaru, 2024; Taichi *et al*., unpublished data). However, it is still unclear why polyploidy enables Cc to be more adaptive to such conditions compared to Ccfc with lower ploidy. Recently, it has been reported that autopolyploidy might contribute to adapting to the urban environment due to the higher reproductive success compared to diploids in *Libidibia ferrea* (Oliveira *et al*., 2022). Autotetraploids of *L. ferrea* have larger flower sizes, likely leading to attracting more pollinators. Autogamous Cc and Ccfc have quite similar flower size, shape, and flowering phenology (Katsuhara and Ushimaru, 2019), thus Cc seems not to have any advantages in these traits compared to Ccfc, unlike the autopolyploids of *L. ferrea*. Allopolyploidy may have a different mechanism leading to obtaining benefits in urban environments for Cc. It is necessary to investigate further details for the relationship between polyploidy and stress tolerance in urban environments.

### Stomata regulated by genetic factors rather than environments

The stomata size and density of Cc and Ccfc (also pollen size, Supplementary data Table S13) appear to be regulated primarily by genetic factors such as genome size, which is consistent with the general finding that genome size correlates positively with stomata size and negatively with stomata density (Beaulieu *et al*., 2008). This idea is also supported by previous studies suggesting that final cell size is only controlled by the minimum set by DNA content, with the results that about 60% of the variation of stomata size (guard cell length) and epidermal cell areas were explained by genome size (Beauliu *et al*., 2008). Considering that Cc was polyploidy and dominant in urban areas, we hypothesize that larger size and lower density of stomata due to polyploidization enhance stress tolerances in urban environments compared to Ccfc. Stomatal morphological traits can be sensitive to environmental changes (Zheng *et al*., 2015) and *Arabidopsis* autotetraploids had larger stomata size and exhibited higher stomatal closure in drought conditions than its diploids, leading to reduced transpiration rate (del Pozo and Ramirez-Parra, 2014). Lower stomata density is also likely to reduce transpiration rates, significantly improving water use efficiency (reviewed by Tossi *et al*., 2022). Thus, larger size and lower density of Cc stomata are likely to be beneficial in stressful environments such as urban drought conditions. However, there were no significant variations in stomata size or density along urban–rural gradients in Cc. Rather, there were some significant differences in these traits due to other factors such as regions, leaf size, and collection dates (Supplementary data Table S11, S12; Fig. S3). In addition, smaller and higher density of stomata are reported to give benefits to faster aperture response (summarized in Bertolino *et al*., 2019). Autotetraploids of other woody species, *Citrus limonia*, exhibited higher drought tolerance with higher ABA content by polyploidization than the diploids, although there were no significant differences in stomata size and density between them (Allario *et al*., 2013). Considering that stomata closure in *Arabidopsis* is controlled by abscisic acid (ABA) and reactive oxygen species (ROS) responses (del Pozo and Ramirez-Parra, 2014), stomata size and density are assumed to indirectly increase the adaptive potential and some functional changes associated with polyploidization may contribute to the improvement of transpiration rate leading to adaptation to urban environments. A respiration and photosynthesis metabolism by changed stomata traits may contribute to expansion in urban environments. Further data needs to be collected and analyzed in combination with stomata traits and other factors to confirm the polyploidy–urban adaptation relationship.

### Scenarios of speciation in Cc, Ccfc, and Cb

Cc (2n = 88) and Ccfc (2n = 46) have genetic variations in chromosome number; as described above and in this study (Fukumoto, 1979; Fujishima, 1981, 2003, 2010; Fig. 4), while Cb is only 2n = 22 in Japan (Fujishima, 2007). Although Cc and Ccfc share several similar morphological traits, previous reports and this study demonstrate that the two species are quite distinct in some aspects of both ecological and genetic traits. This is especially supported by the fact that Cc and Ccfc are seldom fertilized: no seed set was generated between Cc and Ccfc cross (Fujishima, 2003, 2010) and heterospecific pollination produced 11 % of seed sets (Katsuhara and Ushimaru, 2019). As illustrated in Fig. 5 and Supplementary data Fig. S4, the chromosomes of the two species are different in size and structure. Additionally, the GISH image of Cc shows the Ccfc genome with clear fluorescent signals and another unknown genome with no clear signals (Fig. 6). This suggests that Ccfc and another genome may have many similarities. Lee *et al*. (2017) conducted a phylogenetic analysis of the genus *Commelina* in eastern and southeastern Asia. It aims to clarify the taxonomic placement of these species within the family Commelinaceae. A detailed phylogenetic analysis integrating the dataset of plastid genomes such as *matK*, *rbcL*, and ITS regions can be assumed to be a candidate for the donor of the missing genome (Lee *et al*. 2022). In the phylogenetic tree, Cc and Ccfc can be classified as the most closely related sister taxa in the genus *Commelina*. Furthermore, there is also a high degree of overlap between Cc and Ccfc habitats, which is an important factor in the formation of hybrids. There have been no observations of other *Commelina* species which overlap in habitat. Taking account of the phylogenetic analysis and the ecological distribution, one of the potential donors of Cc is likely to be Ccfc. Analysis of microsatellite markers by Katsuhara *et al*. (2019, Table S1) shows that 2 out of 15 markers derived from Ccfc were not amplified in Cc. It suggests that Cc might not have a complete Ccfc chromosome set. Therefore, there is an alternative scenario that Cc and Ccfc share a common ancestor and speciated from that species. In conclusion, Cc has homoeologous chromosomes with Ccfc and Ccfc is the most likely to be one of the donors of Cc. However, it is still unknown which chromosome number is the origin of Cc and Ccfc. Although 2n = 4x = 44 (Ccfc) is likely the origin, there is a problem with dysploidy. In general, dysploidy has two patterns; descending and ascending dysploidy, and descending dysploidy is found in most terrestrial plants while ascending dysploidy is quite rare (Mayrose and Lysak, 2020). It is therefore difficult to say that chromosome numbers are diverse from 2n = 4x = 44 species due to ascending dysploidy. Furthermore, the genus *Commelina* may not have a common basic chromosome number. Cb chromosome number has been reported: 2n = 22, 44, and 66, suggesting the basic number is x = 11 (Fujishima, 2010). However, closely related subspecies, *Commelina diffusa* has different chromosome numbers: *C. diffusa* has 2n = 30, 48, 60 in Thailand, and 2n = 72 in Japan (Fujishima, 2007b; Saensouk and Saensouk, 2020), suggesting basic chromosome number of *C. diffusa* is not x = 11. Thus, ploidy levels and the origin of Cc and Ccfc are still unclear.

Considering Cb and the two species, chromosome size, number of chromosomes, and genome size are quite different. Phylogenetically, Cb is located in different clades from Cc by analysis of plastid *matK* gene (Lee *et al*., 2017). Based on the above genome size and chromosome number, differences in chromosome size might reflect the genome size per chromosome; the genome size per chromosome was estimated to be approximately 160 Mb (Cc), 130 Mb (Ccfc), and 210 Mb (Cb), calculated by dividing the total genome size (1C) by haploid chromosome number. Cc chromosomes had more chromosome knobs and an elongated shape compared to Ccfc and Cb (Fig. 5, Supplementary data S4, S5D). In general, chromosomal knobs are condensed regions of chromatin, heterochromatin where gene expressions are suppressed and they are a chromosomal trait (Fukui *et al*., 1988; Ijima *et al*., 1991). The GISH results suggest the differences in repetitive sequences possession among the three species. For example, Ccfc genomic DNA probe exhibited band signals on Cb chromosomes, which indicates Cb has many amplified specific repetitive sequences (repeats) compared to Ccfc (Fig. 6I). On the other hand, Cb DNA probe appeared dot signals on several Ccfc chromosomes, indicating Ccfc may not have shared repeats as much as Cb (Supplementary data Fig. S5C). When Cb DNA probe was applied to Cc chromosomes, the signal appeared weakly on all chromosomes, suggesting Cc and Cb have less similarity in genomes than Ccfc and Cb (Supplementary data Fig. S5F). A part of the Cc genome is thought to be composed of a different genome from Cb and Ccfc, suggesting the *Commelina* species have a unique and complicated strategy for their diversity. To understand the differences among them, further molecular cytological studies such as repeat analysis are needed.

## CONCLUSIONS

In this study, we focused on *Commelina* species, Cc and its closely related Ccfc, which have high diversity in morphological traits and chromosome number. Our results indicate that Cc is an allopolyploid and we conclude that allopolyploidization might give Cc to have a high potential to survive in urban environments compared to Ccfc. This suggestion is consistent with the fact that Cc populations are spreading at a global scale. Our focus on plant distributions at more microecological scales such as urban– rural gradients demonstrates that environmental changes due to urbanization reveal differences in plant viability particularly in relation to polyploidy. Although pollen fertilities of both Cc and Ccfc are quite high (approx. 90%: unpublished data), Cc and Ccfc seldom cross and generate hybrid seeds (Katsuhara and Ushimaru, 2019), suggesting that they may exhibit difficulties in chromosome pairing in meiotic phases. Further cytogenetic studies such as using the GISH method with other relative species and details of chromosome numbers and chromosome pairing, sequencing, and survival rate experiments are necessary to clarify the polyploidization history which enhances survival potentials of *C. communis* in urban environments. Furthermore, the effects of alloploidy in urban environments need to be verified in other wild plant systems.

