## Supplemental data for "Allopolyploidy enhances survival advantages for urban environments in the native plant genus *Commelina*"

Table S1. Location and distribution ratio data. Site shows the location ID and Site numbers with asterisks indicate that they were excluded data for statistical analysis.

| Site | Region | Localities | Lat (N') | Lon (E') | Distribution ratio | Year |
|---|---|---|---|---|---|---|
| 1 | Shimane | Nakahara, Matsue city, Shimane Pf. | 35°28'07" | 133°02'23" | 5 | 2021 |
| 2 | Shimane | Okudani, Matsue city, Shimane Pf. | 35°28'49" | 133°03'00" | 5 | 2021 |
| 3 | Shimane | Sagusa, Matsue city, Shimane Pf. | 35°25'47" | 133°04'25" | 5 | 2021 |
| 4 | Shimane | Kuroda, Matsue city, Shimane Pf. | 35°28'59" | 133°02'23" | 5 | 2021 |
| 5 | Shimane | Yamashiro, Matsue city, Shimane Pf. | 35°26'39" | 133°05'06" | 5 | 2021 |
| 6 | Shimane | Irie,Yatsuka, Matsue city, Shimane Pf. | 35°29'41" | 133°09'28" | 5 | 2021 |
| 7 | Shimane | Irie,Yatsuka, Matsue city, Shimane Pf. | 35°29'45" | 133°09'31" | 5 | 2021 |
| 8 | Shimane | Osoe,Yatsuka, Matsue city, Shimane Pf. | 35°29'48" | 133°11'15" | 5 | 2021 |
| 9 | Shimane | Ejima,Yatsuka, Matsue city, Shimane Pf. | 35°30'48" | 133°11'34" | 5 | 2021 |
| 10 | Shimane | Ejima,Yatsuka, Matsue city, Shimane Pf. | 35°30'46" | 133°12'01" | 5 | 2021 |
| 11 | Shimane | Kuroda, Matsue city, Shimane Pf. | 35°28'48" | 133°02'48" | 5 | 2021 |
| 12 | Shimane | Tono, Matsue city, Shimane Pf. | 35°28'35" | 133°03'02" | 5 | 2021 |
| 13 | Shimane | Sadahongo, Kashima, Matsue city, Shimane Pf. | 35°31'22" | 132°59'31" | 5 | 2021 |
| 14 | Shimane | Koura, Kashima, Matsue city, Shimane Pf. | 35°31'11" | 132°58'32" | 5 | 2021 |
| 15 | Shimane | Tayu, Kashima, Matsue city, Shimane Pf. | 35°32'12" | 132°58'21" | 5 | 2021 |
| 16 | Shimane | Kataku, Kashima, Matsue city, Shimane Pf. | 35°32'33" | 132°58'58" | 5 | 2021 |
| 17 | Shimane | Mitsu, Kashima, Matsue city, Shimane Pf. | 35°32'24" | 133°01'29" | 5 | 2021 |
| 18 | Shimane | Kitakoubu, Kashima, Matsue city, Shimane Pf. | 35°31'28" | 133°01'29" | 5 | 2021 |
| 19 | Shimane | Kamiyamasa, Hirose, Yasugi city, Shimane Pf. | 35°20'29" | 133°08'08" | 4 | 2021 |
| 20 | Shimane | Kamiyamasa, Hirose, Yasugi city, Shimane Pf. | 35°20'35" | 133°08'04" | 2 | 2021 |

| | | | | | | |
|---|---|---|---|---|---|---|
| 21 | Shimane | Kamiito, Higashiizumo, Matsue city, Shimane Pf. | 35°24'16" | 133°09'23" | 5 | 2021 |
| 22 | Shimane | Kamiito, Higashiizumo, Matsue city, Shimane Pf. | 35°25'13" | 133°09'54" | 5 | 2021 |
| 23 | Shimane | Fukutomi, Matsue city, Shimane Pf. | 35°27'17" | 133°07'27" | 5 | 2021 |
| 24 | Shimane | Fukutomi, Matsue city, Shimane Pf. | 35°27'20" | 133°07'13" | 5 | 2021 |
| 25 | Shimane | Kumano,Yakumo, Matsue city, Shimane Pf. | 35°22'24" | 133°04'16" | 5 | 2021 |
| 26 | Shimane | Kumano,Yakumo, Matsue city, Shimane Pf. | 35°22'23" | 133°04'16" | 5 | 2021 |
| 27 | Shimane | Nishiiwasaka,Yakumo, Matsue city, Shimane Pf. | 35°23'20" | 133°06'04" | 5 | 2021 |
| 28 | Shimane | Nishiiwasaka,Yakumo, Matsue city, Shimane Pf. | 35°24'21" | 133°05'31" | 5 | 2021 |
| 29 | Shimane | Asakakumi, Matsue city, Shimane Pf. | 35°28'27" | 133°06'36" | 5 | 2021 |
| 30 | Shimane | Kamiubeo, Matsue city, Shimane Pf. | 35°29'15" | 133°07'32" | 5 | 2021 |
| 31 | Shimane | Shimokuno, Daito, Unnan city, Shimane Pf. | 35°15'42" | 132°58'55" | 2 | 2022 |
| 32 | Shimane | Kamikuno, Daito, Unnan city, Shimane Pf. | 35°16'03" | 133°02'11" | 5 | 2022 |
| 33 | Shimane | Sajiro, Okuizumo, Nita-gun, Shimane Pf. | 35°13'33" | 132°58'41" | 4 | 2022 |
| 34 | Shimane | Otani, Okuizumo, Nita-gun, Shimane Pf. | 35°10'29" | 133°02'20" | 4 | 2022 |
| 35 | Shimane | Yokota, Okuizumo, Nita-gun, Shimane Pf. | 35°11'16" | 133°04'50" | 5 | 2022 |
| 36 | Shimane | Nishihida, Hirose, Yasugi city, Shimane Pf. | 35°14'18" | 133°07'55" | 2 | 2022 |
| 37 | Shimane | Hube, Hirose, Yasugi city, Shimane Pf. | 35°18'31" | 133°09'15" | 5 | 2022 |
| 38 | Shimane | Sugahara, Hirose, Yasugi city, Shimane Pf. | 35°19'44" | 133°10'06" | 5 | 2022 |
| 39 | Shimane | Higashiiwasaka, Yakumo, Matsue city, Shimane Pf. | 35°22'51" | 133°07'56" | 5 | 2022 |
| 40 | Shimane | Inahara, Okuizumo, Nita-gun, Shimane Pf. | 35°10'06" | 133°06'23" | 4 | 2022 |
| 41 | Shimane | Misawa, Okuizumo, Nita-gun, Shimane Pf. | 35°12'18" | 132°58'45" | 4 | 2022 |
| 42 | Okayama | Tsushimanaka, Kita, Okayama city, Okayama Pf. | 34°41'16" | 133°55'26" | 5 | 2021 |
| 43 | Okayama | Tsushimanaka, Kita, Okayama city, Okayama Pf. | 34°41'06" | 133°55'05" | 5 | 2021 |

| | | | | | | |
|---|---|---|---|---|---|---|
| 44 | Okayama | Houkaiin, Kita, Okayama city, Okayama Pf. | 34°41'32" | 133°56'03" | 5 | 2021 |
| 45 | Okayama | Izumi, Kita, Okayama city, Okayama Pf. | 34°40'44" | 133°55'09" | 5 | 2021 |
| 46 | Okayama | Gakunan, Kita, Okayama city, Okayama Pf. | 34°40'45" | 133°55'27" | 1 | 2021 |
| 47 | Okayama | Minamigata, Kita, Okayama city, Okayama Pf. | 34°40'20" | 133°55'28" | 5 | 2021 |
| 48 | Okayama | R.Nishi,Yanagi, Kita, Okayama city, Okayama Pf. | 34°39'24" | 133°55'22" | 5 | 2021 |
| 49 | Okayama | Noda, Kita, Okayama city, Okayama Pf. | 34°39'15" | 133°54'21" | 5 | 2021 |
| 50 | Okayama | Ima, Kita, Okayama city, Okayama Pf. | 34°38'53" | 133°53'39" | 5 | 2021 |
| 51 | Okayama | R.Sasase, Nishinagase, Kita, Okayama city, Okayama Pf. | 34°38'54" | 133°52'39" | 5 | 2021 |
| 52 | Okayama | Nodonishi, Kita, Okayama city, Okayama Pf. | 34°39'44" | 133°52'50" | 5 | 2021 |
| 53 | Okayama | Nishinoyama, Kita, Okayama city, Okayama Pf. | 34°40'03" | 133°52'42" | 5 | 2021 |
| 54 | Okayama | R.Sasase, Narazu, Kita, Okayama city, Okayama Pf. | 34°40'46" | 133°53'30" | 5 | 2021 |
| 55 | Okayama | Daianjihigashi, Kita, Okayama city, Okayama Pf. | 34°39'53" | 133°53'36" | 5 | 2021 |
| 56 | Okayama | Higashihurumatsu, Kita, Okayama city, Okayama Pf. | 34°38'55" | 133°54'55" | 5 | 2021 |
| 57 | Okayama | Mitsunakayama, Kita, Okayama city, Okayama Pf. | 34°45'31" | 133°55'34" | 5 | 2021 |
| 58 | Okayama | Mitsukusou, Kita, Okayama city, Okayama Pf. | 34°48'53" | 133°56'26" | 5 | 2021 |
| 59 | Okayama | Mitsukusou, Kita, Okayama city, Okayama Pf. | 34°49'37" | 133°55'13" | 5 | 2021 |
| 60 | Okayama | Oda, Tatebe, Kita, Okayama city, Okayama Pf. | 34°51'58" | 133°55'52" | 5 | 2021 |
| 61 | Okayama | Oda, Tatebe, Kita, Okayama city, Okayama Pf. | 34°51'46" | 133°55'59" | 4 | 2021 |
| 62 | Okayama | Oda, Tatebe, Kita, Okayama city, Okayama Pf. | 34°52'04" | 133°57'21" | 4 | 2021 |
| 63 | Okayama | Burakuji, Tatebe, Kita, Okayama city, Okayama Pf. | 34°52'47" | 133°54'47" | 4 | 2021 |
| 64 | Okayama | Shimokoume, Tatebe, Kita, Okayama city, Okayama Pf. | 34°52'53" | 133°55'09" | 3 | 2021 |
| 65 | Okayama | Nishihara, Tatebe, Kita, Okayama city, Okayama Pf. | 34°49'55" | 133°54'05" | 4 | 2021 |
| 66 | Okayama | Mitsutakazu, Kita, Okayama city, Okayama Pf. | 34°48'27" | 133°53'42" | 5 | 2021 |

| | | | | | | |
|---|---|---|---|---|---|---|
| 67 | Okayama | Mitsutakouchi, Kita, Okayama city, Okayama Pf. | 34°46'42" | 133°53'51" | 4 | 2021 |
| 68 | Okayama | Sugano, Kita, Okayama city, Okayama Pf. | 34°45'14" | 133°52'27" | 2 | 2021 |
| 69 | Okayama | Sugano, Kita, Okayama city, Okayama Pf. | 34°44'56" | 133°53'12" | 1 | 2021 |
| 70 | Okayama | Tabara, Kita, Okayama city, Okayama Pf. | 34°44'39" | 133°52'31" | 5 | 2021 |
| 71 | Okayama | Yamanoue, Kita, Okayama city, Okayama Pf. | 34°46'44" | 133°50'28" | 5 | 2021 |
| 72 | Hyogo | Rokkodai, Nada, Kobe city, Hyogo Pf. | 34°43'47" | 135°14'03" | 5 | 2021 |
| 73 | Hyogo | Turukabuto, Nada, Kobe city, Hyogo Pf. | 34°43'47" | 135°14'13" | 5 | 2021 |
| 74 | Hyogo | Mikage, Higashinada, Kobe city, Hyogo Pf. | 34°43'26" | 135°15'20" | 5 | 2021 |
| 75 | Hyogo | R.Toga, Nada, Kobe city, Hyogo Pf. | 34°42'47" | 135°13'44" | 5 | 2021 |
| 76 | Hyogo | Sumadera, Suma, Kobe city, Hyogo Pf. | 34°38'59" | 135°06'47" | 5 | 2021 |
| 77 | Hyogo | Sumadera, Suma, Kobe city, Hyogo Pf. | 34°38'57" | 135°06'42" | 5 | 2021 |
| 78 | Hyogo | Miyuki, Suma, Kobe city, Hyogo Pf. | 34°38'44" | 135°07'18" | 5 | 2021 |
| 79 | Hyogo | Suma seaside park, Suma, Kobe city, Hyogo Pf. | 34°38'38" | 135°07'24" | 5 | 2021 |
| 80* | Hyogo | Suma seaside park, Suma, Kobe city, Hyogo Pf. | 34°38'35" | 135°07'31" | 0 | 2021 |
| 81* | Hyogo | Port Island, Tyuou, Kobe city, Hyogo Pf. | 34°39'41" | 135°13'00" | 0 | 2021 |
| 82 | Hyogo | Port Island, Tyuou, Kobe city, Hyogo Pf. | 34°39'46" | 135°12'53" | 5 | 2021 |
| 83 | Hyogo | Fukae, Higashinada, Kobe city, Hyogo Pf. | 34°43'07" | 135°17'29" | 5 | 2021 |
| 84 | Hyogo | Rokkosan, Nada, Kobe city, Hyogo Pf. | 34°44'39" | 135°11'05" | 5 | 2021 |
| 85 | Hyogo | Rokkosan, Nada, Kobe city, Hyogo Pf. | 34°45'01" | 135°12'41" | 5 | 2021 |
| 86 | Hyogo | Rokkosan, Nada, Kobe city, Hyogo Pf. | 34°45'43" | 135°14'23" | 5 | 2021 |
| 87 | Hyogo | Rokkosan, Nada, Kobe city, Hyogo Pf. | 34°45'48" | 135°14'31" | 5 | 2021 |
| 88 | Hyogo | Kamiyamaguchi, Yamaguchi, Nishinomiya city, Hyogo Pf. | 34°49'17" | 135°14'33" | 2 | 2021 |
| 89 | Hyogo | Kamiyamaguchi, Yamaguchi, Nishinomiya city, Hyogo Pf. | 34°49'18" | 135°14'42" | 4 | 2021 |

| | | | | | | |
|---|---|---|---|---|---|---|
| 90 | Hyogo | Shiose, Nishinomiya city, Hyogo Pf. | 34°50'26" | 135°15'48" | 3 | 2021 |
| 91 | Hyogo | Shiose, Nishinomiya city, Hyogo Pf. | 34°50'29" | 135°15'51" | 2 | 2021 |
| 92 | Hyogo | Minamigaoka, Sanda city, Hyogo Pf. | 34°52'50" | 135°13'20" | 3 | 2021 |
| 93 | Hyogo | Sumada, Sanda city, Hyogo Pf. | 34°57'29" | 135°11'54" | 4 | 2021 |
| 94 | Hyogo | Sumada, Sanda city, Hyogo Pf. | 34°57'28" | 135°11'52" | 2 | 2021 |
| 95 | Hyogo | Sumada, Sanda city, Hyogo Pf. | 34°57'25" | 135°11'50" | 3 | 2021 |
| 96 | Hyogo | Shimosasori, Takarazuka city, Hyogo Pf. | 34°55'35" | 135°18'53" | 4 | 2021 |
| 97 | Hyogo | Shimosasori, Takarazuka city, Hyogo Pf. | 34°55'32" | 135°18'52" | 5 | 2021 |
| 98 | Hyogo | Tamida, Inagawa, Kawabe county, Hyogo Pf. | 34°56'01" | 135°23'35" | 3 | 2021 |
| 99 | Hyogo | Tamida, Inagawa, Kawabe county, Hyogo Pf. | 34°56'03" | 135°23'31" | 3 | 2021 |
| 100 | Hyogo | Arino, Kita, Kobe city, Hyogo Pf. | 34°47'24" | 135°13'10" | 4 | 2021 |
| 101 | Hyogo | Yata, Kita, Kobe city, Hyogo Pf. | 34°49'19" | 135°10'56" | 5 | 2021 |
| 102 | Hyogo | Yata, Kita, Kobe city, Hyogo Pf. | 34°49'20" | 135°11'01" | 5 | 2021 |
| 103 | Hyogo | Oshibedani, Nishi-ku, Kobe city, Hyogo Pf. | 34°44'18" | 135°02'14" | 5 | 2022 |
| 104 | Hyogo | Oshibedani, Nishi-ku, Kobe city, Hyogo Pf. | 34°44'22" | 135°04'24" | 2 | 2022 |
| 105 | Hyogo | Igawadani, Nishi-ku, Kobe city, Hyogo Pf. | 34°41'33" | 135°03'12" | 5 | 2022 |
| 106 | Hyogo | Hasetani, Nishi-ku, Kobe city, Hyogo Pf. | 34°42'33" | 135°01'12" | 5 | 2022 |
| 107 | Hyogo | Shimotanigami, Yamada, Kita-ku, Kobe city, Hyogo Pf. | 34°45'37" | 135°09'16" | 4 | 2022 |
| 108 | Hyogo | Aina, Yamada, Kita-ku, Kobe city, Hyogo Pf. | 34°43'39" | 135°06'41" | 4 | 2022 |
| 109 | Hyogo | Nishishimo, Yamada, Kita-ku, Kobe city, Hyogo Pf. | 34°45'38" | 135°06'46" | 3 | 2022 |
| 110 | Hyogo | Shijimi, Miki city, Hyogo Pf. | 34°46'55" | 135°04'06" | 4 | 2022 |
| 111 | Hyogo | Ogo, Kita-ku, Kobe city, Hyogo Pf. | 34°48'36" | 135°05'48" | 4 | 2022 |
| 112 | Hyogo | Yata, Kita-ku, Kobe city, Hyogo Pf. | 34°49'19" | 135°10'56" | 4 | 2022 |

| | | | | | | |
|---|---|---|---|---|---|---|
| 113 | Hyogo | Arino, Kita-ku, Kobe city, Hyogo Pf. | 34°46'40" | 135°12'16" | 3 | 2022 |
| 114 | Osaka | Yono,Toyono,Toyono county, Osaka Pf. | 34°54'60" | 135°29'37" | 1 | 2021 |
| 115 | Osaka | Yono,Toyono,Toyono county, Osaka Pf. | 34°54'58" | 135°29'31" | 1 | 2021 |
| 116 | Osaka | Izumihara, Ibaraki city, Osaka Pf. | 34°52'53" | 135°31'26" | 5 | 2021 |
| 117 | Osaka | R. saho, Izumihara, Ibaraki city, Osaka Pf. | 34°53'05" | 135°31'39" | 4 | 2021 |
| 118 | Osaka | Izumihara, Ibaraki city, Osaka Pf. | 34°53'03" | 135°31'31" | 1 | 2021 |
| 119 | Osaka | Ichihara, Takatsuki city, Osaka Pf. | 34°55'34" | 135°35'56" | 5 | 2021 |
| 120 | Osaka | Ichihara, Takatsuki city, Osaka Pf. | 34°54'30" | 135°36'08" | 5 | 2021 |
| 121 | Osaka | Ichihara, Takatsuki city, Osaka Pf. | 34°53'24" | 135°36'02" | 2 | 2021 |
| 122 | Osaka | Ichihara, Takatsuki city, Osaka Pf. | 34°53'23" | 135°35'53" | 5 | 2021 |
| 123 | Osaka | Ozato, Nose,Toyono county, Osaka Pf. | 34°58'25" | 135°24'42" | 3 | 2021 |
| 124 | Osaka | Hirano, Nose,Toyono county, Osaka Pf. | 34°57'16" | 135°24'17" | 2 | 2021 |
| 125 | Osaka | Shimoda, Nose,Toyono county, Osaka Pf. | 34°57'13" | 135°24'23" | 5 | 2021 |
| 126 | Osaka | Gein, Mino city, Osaka Pf. | 34°50'15" | 135°30'02" | 4 | 2021 |
| 127 | Osaka | Fujishirodai, Suita city, Osaka Pf. | 34°49'03" | 135°30'46" | 5 | 2021 |
| 128 | Osaka | Yamadakita, Suita city, Osaka Pf. | 34°48'31" | 135°31'06" | 5 | 2021 |
| 129 | Osaka | Tsukumodai, Suita city, Osaka Pf. | 34°48'05" | 135°30'53" | 5 | 2021 |
| 130 | Osaka | Tsukumodai, Suita city, Osaka Pf. | 34°47'48" | 135°30'36" | 5 | 2021 |
| 131 | Osaka | Senriyamatakezono, Suita city, Osaka Pf. | 34°47'10" | 135°30'05" | 2 | 2021 |
| 132 | Osaka | Kasuga, Suita city, Osaka Pf. | 34°47'09" | 135°29'57" | 5 | 2021 |
| 133 | Osaka | Hattoriryokuchi, Toyonaka city, Osaka Pf. | 34°46'45" | 135°29'41" | 5 | 2021 |
| 134 | Osaka | Nishiizumigaoka, Toyonaka city, Osaka Pf. | 34°46'54" | 135°29'12" | 5 | 2021 |
| 135 | Osaka | Nishiizumigaoka, Toyonaka city, Osaka Pf. | 34°47'06" | 135°29'12" | 5 | 2021 |

| | | | | | | |
|---|---|---|---|---|---|---|
| 136 | Osaka | Senribanbaku park, Suita city, Osaka Pf. | 34°48'34" | 135°31'55" | 5 | 2021 |
| 137 | Osaka | Senribanbaku park, Suita city, Osaka Pf. | 34°48'38" | 135°32'12" | 5 | 2021 |
| 138 | Osaka | Minato,Naniwa, Osaka city, Osaka Pf. | 34°40'08" | 135°29'47" | 5 | 2021 |
| 139 | Osaka | Shinmachi, Nishi, Osaka city, Osaka Pf. | 34°40'38" | 135°29'27" | 5 | 2021 |
| 140 | Osaka | Nihonbashi, Chuou, Osaka city, Osaka Pf. | 34°40'01" | 135°30'25" | 5 | 2021 |
| 141 | Osaka | Nihonbashi, Chuou, Osaka city, Osaka Pf. | 34°39'59" | 135°31'00" | 5 | 2021 |
| 142 | Osaka | Obase, Tennoji, Osaka city, Osaka Pf. | 34°39'57" | 135°31'31" | 5 | 2021 |
| 143 | Osaka | Kitahamahigashi, Chuou, Osaka city, Osaka Pf. | 34°41'23" | 135°30'37" | 5 | 2021 |
| 144 | Osaka | Minamiterakatahigashi, Moriguchi city, Osaka Pf. | 34°43'02" | 135°34'33" | 5 | 2021 |
| 145 | Osaka | Shodainakamachi, Hirakata city, Osaka Pf. | 34°50'20" | 135°41'17" | 5 | 2022 |
| 146 | Osaka | Kasugahigashi, Hirakata city, Osaka Pf. | 34°48'15" | 135°41'36" | 5 | 2022 |
| 147 | Osaka | Sonenji, Hirakata city, Osaka Pf. | 34°47'55" | 135°43'29" | 4 | 2022 |
| 148* | Osaka | Mitsuya, Hirakata city, Osaka Pf. | 34°49'02" | 135°38'30" | 0 | 2022 |
| 149* | Osaka | Higashiamagawa, Takatsuki city, Osaka Pf. | 34°50'56" | 135°38'38" | 0 | 2022 |
| 150 | Osaka | Karasakinaka, Takatsuki city, Osaka Pf. | 34°48'42" | 135°36'59" | 5 | 2022 |
| 151 | Kyoto | Fukakusasengahara, Fushimi, Kyoto city, Kyoto Pf. | 34°57'39" | 135°47'31" | 4 | 2021 |
| 152 | Kyoto | Kanshujidainichi, Yamashina, Kyoto city, Kyoto Pf. | 34°57'32" | 135°47'35" | 3 | 2021 |
| 153 | Kyoto | Sagakankujikubodono, Ukyo, Kyoto city, Kyoto Pf. | 35°01'36" | 135°40'30" | 4 | 2021 |
| 154 | Kyoto | Sagakankujikubodono, Ukyo, Kyoto city, Kyoto Pf. | 35°01'38" | 135°40'34" | 5 | 2021 |
| 155 | Kyoto | Oharanokitakasuga, Saikyo, Kyoto city, Kyoto Pf. | 34°57'56" | 135°39'35" | 5 | 2021 |
| 156 | Kyoto | Kyotogyoen, Kamikyo, Kyoto city, Kyoto Pf. | 35°01'38" | 135°45'36" | 5 | 2021 |
| 157 | Kyoto | Kyotogyoen, Kamikyo, Kyoto city, Kyoto Pf. | 35°01'10" | 135°45'35" | 5 | 2021 |
| 158 | Kyoto | Nishikyogokukitashozakai, Ukyo, Kyoto city, Kyoto Pf. | 34°59'33" | 135°43'37" | 5 | 2021 |

| | | | | | | |
|---|---|---|---|---|---|---|
| 159 | Kyoto | Kamitobanishiura, Minami, Kyoto city, Kyoto Pf. | 34°57'25" | 135°44'25" | 5 | 2021 |
| 160 | Kyoto | Mukoujimafujinoki, Fushimi, Kyoto city, Kyoto Pf. | 34°55'30" | 135°46'50" | 5 | 2021 |
| 161 | Kyoto | Mukoujimafujinoki, Fushimi, Kyoto city, Kyoto Pf. | 34°55'25" | 135°46'47" | 5 | 2021 |
| 162 | Kyoto | Nijojo, Tyukyo, Kyoto city, Kyoto Pf. | 35°00'45" | 135°44'54" | 5 | 2021 |
| 163 | Kyoto | Maruyama, Higashiyama, Kyoto city, Kyoto Pf. | 35°00'13" | 135°46'49" | 5 | 2021 |
| 164 | Kyoto | Kankiji, Shimokyo, Kyoto city, Kyoto Pf. | 34°59'11" | 135°44'44" | 5 | 2021 |
| 165 | Kyoto | Yokoohjikuwanomoto, Fushimi, Kyoto city, Kyoto Pf. | 34°55'30" | 135°44'20" | 5 | 2021 |
| 166 | Kyoto | Yamanouchimiyawaki, Ukyo, Kyoto city, Kyoto Pf. | 35°00'41" | 135°43'27" | 5 | 2021 |
| 167 | Kyoto | Nishigamo, Kita, Kyoto city, Kyoto Pf. | 35°03'51" | 135°44'34" | 5 | 2021 |
| 168 | Kyoto | Shizuichishizuhara, Sakyo, Kyoto city, Kyoto Pf. | 35°04'21" | 135°45'59" | 4 | 2021 |
| 169 | Kyoto | Iwakura, Sakyo, Kyoto city, Kyoto Pf. | 35°05'34" | 135°47'22" | 4 | 2021 |
| 170 | Kyoto | Yase, Sakyo, Kyoto city, Kyoto Pf. | 35°04'60" | 135°49'06" | 5 | 2021 |
| 171 | Kyoto | Ohara, Sakyo, Kyoto city, Kyoto Pf. | 35°08'41" | 135°50'22" | 2 | 2021 |
| 172 | Kyoto | Momoi, Ohara, Sakyo, Kyoto city, Kyoto Pf. | 35°09'44" | 135°48'35" | 2 | 2021 |
| 173 | Kyoto | Momoi, Ohara, Sakyo, Kyoto city, Kyoto Pf. | 35°09'45" | 135°48'30" | 2 | 2021 |
| 174 | Kyoto | Hanase, Sakyo, Kyoto city, Kyoto Pf. | 35°09'54" | 135°46'50" | 3 | 2021 |
| 175 | Kyoto | Hanase, Sakyo, Kyoto city, Kyoto Pf. | 35°09'53" | 135°46'59" | 3 | 2021 |
| 176 | Kyoto | Hanase, Sakyo, Kyoto city, Kyoto Pf. | 35°09'60" | 135°46'51" | 4 | 2021 |
| 177 | Kyoto | Shizuichiichihara, Sakyo, Kyoto city, Kyoto Pf. | 35°06'27" | 135°47'09" | 2 | 2021 |
| 178 | Kyoto | Shizuichiichihara, Sakyo, Kyoto city, Kyoto Pf. | 35°04'54" | 135°45'36" | 2 | 2021 |
| 179 | Kyoto | Dannomachi, Kamiueno, Mukou city, Kyoto Pf. | 34°55'49" | 135°42'09" | 5 | 2022 |
| 180 | Kyoto | Yodomizu, Fushimi-ku, Kyoto city, Kyoto Pf. | 34°53'49" | 135°42'32" | 5 | 2022 |
| 181 | Kyoto | Mukaijimanishi, Fushimi-ku, Kyoto city, Kyoto Pf. | 34°54'34" | 135°45'51" | 5 | 2022 |

| | | | | | | |
|---|---|---|---|---|---|---|
| 182 | Kyoto | Kumiyama, Kuse-gun, Kyoto Pf. | 34°52'53" | 135°45'44" | 5 | 2022 |
| 183 | Kyoto | Kumiyama, Kuse-gun, Kyoto Pf. | 34°53'47" | 135°43'43" | 5 | 2022 |
| 184 | Kyoto | Yawata, Yawata city, Kyoto Pf. | 34°52'28" | 135°42'43" | 4 | 2022 |
| 185 | Kyoto | Yawata, Yawata city, Kyoto Pf. | 34°52'40" | 135°41'48" | 5 | 2022 |
| 186* | Kyoto | Yawata, Yawata city, Kyoto Pf. | 34°53'13" | 135°41'57" | 0 | 2022 |
| 187 | Kyoto | Matsuimukai, Kyotanabe city, Kyoto Pf. | 34°50'49" | 135°44'16" | 5 | 2022 |
| 188* | Kyoto | Tanabehigashihama, Kyotanabe city, Kyoto Pf. | 34°49'44" | 135°46'05" | 0 | 2022 |
| 189 | Kyoto | Tonouchikawa, Joyo city, Kyoto Pf. | 34°50'03" | 135°47'07" | 5 | 2022 |
| 190* | Kyoto | Tonotakai, Joyo city, Kyoto Pf. | 34°50'45" | 135°46'22" | 0 | 2022 |
| 191 | Tokyo | Yumenoshima, Koto-ku, Tokyo Pf. | 35°39'04" | 139°49'40" | 5 | 2022 |
| 192 | Tokyo | Hibiyakoen, Chiyoda-ku, Tokyo Pf. | 35°40'29" | 139°45'30" | 5 | 2022 |
| 193 | Tokyo | Kitanomarukoen, Chiyoda-ku, Tokyo Pf. | 35°41'32" | 139°45'02" | 5 | 2022 |
| 194 | Tokyo | Kamizono, Yoyogi, Shibuya-ku, Tokyo Pf. | 35°40'20" | 139°42'01" | 5 | 2022 |
| 195 | Tokyo | Shiroganedai, Minato-ku, Tokyo Pf. | 35°38'11" | 139°43'08" | 2 | 2022 |
| 196 | Tokyo | Heiwanomorikouen, Oda-ku, Tokyo Pf. | 35°34'46" | 139°44'28" | 5 | 2022 |
| 197 | Tokyo | Ikegami, Oda-ku, Tokyo Pf. | 35°34'43" | 139°42'19" | 5 | 2022 |
| 198 | Tokyo | Minamizenzoku, Oda-ku, Tokyo Pf. | 35°36'10" | 139°41'29" | 5 | 2022 |
| 199 | Tokyo | Kinutakoen, Setagaya-ku, Tokyo Pf. | 35°37'41" | 139°37'30" | 5 | 2022 |
| 200 | Tokyo | Wakaba, Chohu city, Tokyo Pf. | 35°39'26" | 139°34'53" | 5 | 2022 |
| 201 | Tokyo | Gotenyama, Musashino city, Tokyo Pf. | 35°42'00" | 139°34'38" | 5 | 2022 |
| 202 | Tokyo | Osawa, Mitaka city, Tokyo Pf. | 35°40'43" | 139°31'51" | 5 | 2022 |
| 203 | Tokyo | Higashi, Koganei city, Tokyo Pf. | 35°41'08" | 139°31'32" | 4 | 2022 |

| | | | | | | |
|---|---|---|---|---|---|---|
| 204 | Tokyo | Osawa, Mitaka city, Tokyo Pf. | 35°41'02" | 139°31'21" | 5 | 2022 |
| 205 | Tokyo | Higashinaganuma, Inagi city, Tokyo Pf. | 35°38'58" | 139°30'13" | 5 | 2022 |
| 206 | Tokyo | Sekido, Tama city, Tokyo Pf. | 35°38'20" | 139°26'52" | 5 | 2022 |
| 207 | Tokyo | Renkoji, Tama city, Tokyo Pf. | 35°38'16" | 139°27'47" | 5 | 2022 |
| 208 | Tokyo | Renkoji, Tama city, Tokyo Pf. | 35°38'13" | 139°27'43" | 4 | 2022 |
| 209 | Tokyo | Taniho, Kunitachi city, Tokyo Pf. | 35°40'37" | 139°26'50" | 5 | 2022 |
| 210 | Tokyo | Kamioyamada, Machida city, Tokyo Pf. | 35°36'23" | 139°23'58" | 5 | 2022 |
| 211 | Tokyo | Kishi, Musashimurayama city, Tokyo Pf. | 35°46'11" | 139°22'02" | 2 | 2022 |
| 212* | Tokyo | Hanekami, Hamura city, Tokyo Pf. | 35°45'37" | 139°18'07" | — | 2022 |
| 213 | Tokyo | Hinode, Tama-gun, Tokyo Pf. | 35°45'46" | 139°13'35" | 5 | 2022 |
| 214 | Tokyo | Sawai, Oume city, Tokyo Pf. | 35°48'30" | 139°11'03" | 5 | 2022 |
| 215 | Tokyo | Okutama, Nishitama-gun, Tokyo Pf. | 35°48'41" | 139°07'24" | 4 | 2022 |
| 216 | Tokyo | Itsukaichi, Akiruno city, Tokyo Pf. | 35°43'37" | 139°13'33" | 3 | 2022 |
| 217 | Tokyo | Hinoharamura, Nishitama-gun, Tokyo Pf. | 35°44'26" | 139°08'44" | 3 | 2022 |
| 218 | Tokyo | Kamiongata, Hachioji city, Tokyo Pf. | 35°40'05" | 139°14'02" | 4 | 2022 |

Table S2. Cultivated samples in 2021 and 2022. Cc and Ccfc indicate *C. communis* and *C. c.* f. *ciliata*, respectively.

| Region | Cc | Ccfc | Cc | Ccfc |
|---|---|---|---|---|
| Tokyo | - | - | 15 | 11 |
| Kyoto | 6 | 6 | 6 | 3 |
| Osaka | 3 | 3 | - | - |
| Hyogo | 4 | 3 | 3 | - |
| Shimane | 14 | 4 | - | - |
| Total | 27 | 16 | 24 | 14 |

Table S3. Location data of materials for stomata analysis. Site numbers are the same as the data shown in Table S1.

| Name | Species | Region | Localities | Lat (N') | Lon (E') |
|---|---|---|---|---|---|
| Sajiro | *C. c.* f. *ciliata* | Shimane | Site 33 | 35°13'33" | 132°58'41" |
| Okuizumo | *C. c.* f. *ciliata* | Shimane | Site 41 | 35°12'18" | 132°58'45" |
| Nunobe | *C. c.* f. *ciliata* | Shimane | Site 37 | 35°18'31" | 133°09'15" |
| Kisuki | *C. communis* | Shimane | Site 31 | 35°15'42" | 132°58'55" |
| Okuizumo | *C. communis* | Shimane | Site 41 | 35°12'18" | 132°58'45" |
| Nunone | *C. communis* | Shimane | Site 37 | 35°18'31" | 133°09'15" |
| Uchinakahara | *C. communis* | Shimane | Uchinakahara, Matsue city, Shimane Pf. | 35°28'23" | 133°02'47" |
| Shimane Univ. | *C. communis* | Shimane | Nishikawatsu, Matsue city, Shimane Pf. | 35°29'07" | 133°04'13" |
| Akayama | *C. communis* | Shimane | Kitahori, Matsue city, Shimane Pf. | 35°28'43" | 133°03'10" |
| Oshibedani | *C. c.* f. *ciliata* | Hyogo | Site 104 | 34°44'22" | 135°04'24" |
| Minotani | *C. c.* f. *ciliata* | Hyogo | Site 107 | 34°45'37" | 135°09'16" |
| Shijimi | *C. c.* f. *ciliata* | Hyogo | Site 110 | 34°46'55" | 135°04'06" |
| Oshibedani | *C. communis* | Hyogo | Site 104 | 34°44'22" | 135°04'24" |
| Minotani | *C. communis* | Hyogo | Site 107 | 34°45'37" | 135°09'16" |
| Shijimi | *C. communis* | Hyogo | Site 110 | 34°46'55" | 135°04'06" |
| Rokkomichi | *C. communis* | Hyogo | Fukada, Nada, Kobe city, Hyogo Pf. | 34°42'56" | 135°14'24" |
| Togagawa | *C. communis* | Hyogo | Shinoharahonmachi, Nada, Kobe city, Hyogo Pf. | 34°43'14" | 135°13'44" |
| Sakuragaoka | *C. communis* | Hyogo | Sakuragaoka, Nada, Kobe city, Hyogo Pf. | 34°43'34" | 135°14'26" |
| Shizuhara | *C. c.* f. *ciliata* | Kyoto | Site 177 | 35°06'27" | 135°47'09" |
| Shizuichi | *C. c.* f. *ciliata* | Kyoto | Site 178 | 35°04'54" | 135°45'36" |
| Ohara | *C. c.* f. *ciliata* | Kyoto | Site 171 | 35°08'41" | 135°50'22" |

| | | | | | |
|---|---|---|---|---|---|
| Shizuhara | *C. communis* | Kyoto | Site 177 | 35°06'27" | 135°47'09" |
| Shizuichi | *C. communis* | Kyoto | Site 178 | 35°04'54" | 135°45'36" |
| Ohara | *C. communis* | Kyoto | Site 171 | 35°08'41" | 135°50'22" |
| Minamiku | *C. communis* | Kyoto | Kamitobanishiura, Minami, Kyoto city, Kyoto Pf. | 34°57'22" | 135°44'22" |
| Nishikyogoku | *C. communis* | Kyoto | Site 158 | 34°59'33" | 135°43'38" |
| Nijojo | *C. communis* | Kyoto | Shuzei, Kamigyo, Kyoto city, Kyoto Pf. | 35°01'02" | 135°44'44" |
| Hinohara | *C. c.* f. *ciliata* | Tokyo | Site 217 | 35°44'26" | 139°08'44" |
| Hachioji | *C. c.* f. *ciliata* | Tokyo | Site 218 | 35°40'05" | 139°14'02" |
| Okutama | *C. c.* f. *ciliata* | Tokyo | Site 215 | 35°48'41" | 139°07'25" |
| Hinohara | *C. communis* | Tokyo | Site 217 | 35°44'26" | 139°08'44" |
| Hachioji | *C. communis* | Tokyo | Site 218 | 35°40'05" | 139°14'02" |
| Okutama | *C. communis* | Tokyo | Site 215 | 35°48'41" | 139°07'25" |
| Senzoku | *C. communis* | Tokyo | Site 198 | 35°36'10" | 139°41'29" |
| Saneatsu | *C. communis* | Tokyo | Site 200 | 35°39'26" | 139°34'53" |
| Hamura | *C. communis* | Tokyo | Site 212 | 35°45'37" | 139°18'07" |

Table S4. Material data for flow cytometry analysis. Site numbers are same as the data shown in Table S1.

| Name | Species | Region | Localities |
|---|---|---|---|
| Okudani3 | *C. communis* | Shimane | Site 2 |
| Uchinakahara | *C. communis* | Shimane | Non-recorded |
| Nishiiwasaka1 | *C. communis* | Shimane | Site 27 |
| Nishiiwasaka2 | *C. communis* | Shimane | Site 28 |
| Sagusa | *C. communis* | Shimane | Site 3 |
| Kuroda1 | *C. communis* | Shimane | Site 4 |
| Kamiubeo | *C. communis* | Shimane | Site 30 |
| Hirose1 | *C. c.* f. *ciliata* | Shimane | Site 19 |
| Hirose1 | *C. c.* f. *ciliata* | Shimane | Site 19 |
| Hirose2 | *C. c.* f. *ciliata* | Shimane | Site 20 |
| Kamigamo | *C. communis* | Kyoto | Site 168 |
| Ohara | *C. communis* | Kyoto | Site 171 |
| Kubodono | *C. communis* | Kyoto | Site 153 or 154 |
| Nijojo | *C. communis* | Kyoto | Site 162 |
| Kamigamo | *C. c.* f. *ciliata* | Kyoto | Site 168 |
| Ohara | *C. c.* f. *ciliata* | Kyoto | Site 171 |
| Kubodono | *C. c.* f. *ciliata* | Kyoto | Site 153 or 154 |
| Fukakusa | *C. benghalensis* | Kyoto | Site 151 or 152 |

Table S5. Generalized linear mixed model (GLMM) dataset for comparison of Cc and Ccfc. The data of Cc in Tokyo was calculated as a standard. The ratio of developed land area: Species demonstrates an interaction term. Boldface indicates significant effects ($P < 0.001$, $P < 0.01$, and $P < 0.05$).

| Range | AIC | Explanatory variable | Coefficients | | | |
|---|---|---|---|---|---|---|
| | | | Estimate | SE | z value | *p* |
| 500 m | 276.7 | Presence/ | | | | |
| | | (Intercept) | 4.032 | 0.775 | 5.203 | **p<0.001** |
| | | The ratio of developed land area | 0.006 | 0.015 | 0.398 | 0.691 |
| | | Species (Ccfc) | -3.378 | 0.643 | -5.250 | **p<0.001** |
| | | Region Kyoto | 0.265 | 0.592 | 0.447 | 0.655 |
| | | Region Osaka | -0.415 | 0.626 | -0.663 | 0.507 |
| | | Region Hyogo | 0.387 | 0.583 | 0.664 | 0.507 |
| | | Region Okayama | -0.650 | 0.640 | -1.015 | 0.310 |
| | | Region Shimane | -1.315 | 0.587 | -2.239 | **p<0.05** |
| | | The ratio of developed land area: Species | -0.044 | 0.016 | -2.694 | **p<0.01** |
| 1000 m | 268.2 | Presence/ | | | | |
| | | (Intercept) | 4.380 | 0.803 | 5.452 | **p<0.001** |
| | | The ratio of developed land area | 0.008 | 0.016 | 0.500 | 0.617 |
| | | Species (Ccfc) | -3.244 | 0.632 | -5.133 | **p<0.001** |
| | | Region Kyoto | -0.038 | 0.624 | -0.061 | 0.952 |
| | | Region Osaka | -0.768 | 0.664 | -1.156 | 0.248 |
| | | Region Hyogo | 0.059 | 0.613 | 0.096 | 0.923 |
| | | Region Okayama | -1.096 | 0.677 | -1.618 | 0.106 |
| | | Region Shimane | -1.772 | 0.630 | -2.813 | **p<0.01** |
| | | The ratio of developed land area: Species | -0.051 | 0.017 | -2.988 | **p<0.01** |

Table S6. Generalized linear mixed model (GLMM) dataset for comparison of Cc and Ccfc. Year represents sampling year, and the data of Tokyo was calculated as a standard. Boldface indicates significant effects (P < 0.001 or P < 0.05).

| Range | AIC | Explanatory variable | Coefficients | | | | Threshold coefficients | | | |
|---|---|---|---|---|---|---|---|---|---|---|
| | | | Estimate | SE | z value | *p* | | Estimate | SE | z value |
| 500 m | 399.71 | Artificial fields | 0.039 | 0.007 | 5.823 | **p<0.001** | 1\|2 | -2.927 | 0.752 | -3.894 |
| | | Year | -0.028 | 0.402 | -0.070 | 0.944 | | | | |
| | | Region Kyoto | -0.435 | 0.644 | -0.676 | 0.499 | 2\|3 | -1.237 | 0.645 | -1.919 |
| | | Region Osaka | -0.586 | 0.698 | -0.839 | 0.402 | | | | |
| | | Region Hyogo | -0.348 | 0.612 | -0.569 | 0.569 | 3\|4 | -0.610 | 0.638 | -0.957 |
| | | Region Okayama | 0.258 | 0.749 | 0.344 | 0.731 | | | | |
| | | Region Shimane | 1.448 | 0.671 | 2.159 | **p<0.05** | 4\|5 | 0.437 | 0.640 | 0.684 |
| 1000 m | 391.67 | Artificial fields | 0.044 | 0.007 | 6.161 | **p<0.001** | 1\|2 | -2.764 | 0.766 | -3.608 |
| | | Year | -0.273 | 0.407 | -0.670 | 0.503 | | | | |
| | | Region Kyoto | -0.290 | 0.663 | -0.437 | 0.662 | 2\|3 | -1.072 | 0.661 | -1.622 |
| | | Region Osaka | -0.541 | 0.718 | -0.753 | 0.451 | | | | |
| | | Region Hyogo | -0.165 | 0.626 | -0.264 | 0.792 | 3\|4 | -0.442 | 0.655 | -0.675 |
| | | Region Okayama | 0.440 | 0.765 | 0.575 | 0.566 | | | | |
| | | Region Shimane | 1.751 | 0.687 | 2.549 | **p<0.05** | 4\|5 | 0.645 | 0.659 | 0.978 |

Table S7. Average stomata size and density in regions. The values of stomata size and density are averages of 90 stomata and 9 photos per region in rural areas, respectively. Different letters behind the data indicate the significant difference in species and regions ($p < 0.05$).

| | Stomata size （µm） | | | |
|---|---|---|---|---|
| Species | Shimane | Hyogo | Kyoto | Tokyo |
| *C. communis* | 54.02±4.76 a | 54.00±3.82 ab | 52.12±4.48 ac | 47.49±6.34 bcd |
| *C. c.* f. *ciliata* | 44.78±3.17 ce | 42.50±5.13 de | 44.72±6.67 de | 40.92±3.42 e |
| | Stomata density （number・mm$^{-2}$） | | | |
| *C. communis* | 41.77±5.94 B | 39.66±5.69 B | 39.48±6.76 B | 53.54±18.25 AB |
| *C. c.* f. *ciliata* | 61.59±8.73 A | 65.39±8.54 A | 48.30±15.17 AB | 61.60±12.58 A |

Table S8. Generalized linear mixed model (GLMM) dataset for comparison of Cc and Ccfc in stomata size and density. Boldface indicates significant effects ($P < 0.001$, $P < 0.01$, and $P < 0.05$). P value with asterisk demonstrates a significant tendency.

| Response variable / Explanatory variable | Estimated coefficient | Standard error | *z* value | *P* |
|---|---|---|---|---|
| Stomata size/ | | | | |
| (Intercept) | 47.598 | 1.656 | 28.736 | **<0.001** |
| CcKyoto | 4.718 | 2.309 | 2.043 | **<0.05** |
| CcHyogo | 6.717 | 2.401 | 2.797 | **<0.01** |
| CcShimane | 6.822 | 2.229 | 3.061 | **<0.01** |
| CcfcTokyo | -6.604 | 0.619 | -10.671 | **<0.001** |
| CcfcKyoto | -2.708 | 2.298 | -1.179 | 0.239 |
| CcfcHyogo | -4.823 | 2.358 | -2.045 | **<0.05** |
| CcfcShimane | -1.273 | 2.205 | -0.577 | 0.564 |
| Leaf area | -0.060 | 0.206 | -0.292 | 0.770 |
| Stomata density/ | | | | |
| (Intercept) | 49.324 | 5.064 | 9.740 | **<0.001** |
| CcKyoto | -17.767 | 6.661 | -2.667 | **<0.01** |
| CcHyogo | -22.520 | 7.980 | -2.822 | **<0.01** |
| CcShimane | -19.537 | 7.142 | -2.735 | **<0.01** |
| CcfcTokyo | 9.238 | 4.407 | 2.096 | **<0.05** |
| CcfcKyoto | -7.755 | 6.483 | -1.196 | 0.232 |
| CcfcHyogo | 5.077 | 7.387 | 0.687 | 0.492 |
| CcfcShimane | 2.018 | 6.877 | 0.293 | 0.769 |
| Leaf area | 2.437 | 1.373 | 1.775 | 0.076* |

Table S9. Multiple Comparisons of Means in stomata size: Tukey Contrasts. The boldface indicates significant effects (P < 0.001). P value with asterisk indicates significant effects (P < 0.01).

| Linear Hypotheses | Estimated coefficient | Standard error | *z* value | *P* |
|---|---|---|---|---|
| CcKyoto - CcTokyo == 0 | 4.718 | 2.309 | 2.043 | 0.361 |
| CcHyogo - CcTokyo == 0 | 6.717 | 2.401 | 2.797 | 0.065* |
| CcShimane - CcTokyo == 0 | 6.822 | 2.229 | 3.061 | **<0.05** |
| CcfcTokyo - CcTokyo == 0 | -6.604 | 0.619 | -10.671 | **<0.001** |
| CcfcKyoto - CcTokyo == 0 | -2.708 | 2.298 | -1.179 | 0.906 |
| CcfcHyogo - CcTokyo == 0 | -4.823 | 2.358 | -2.045 | **0.359** |
| CcfcShimane - CcTokyo == 0 | -1.273 | 2.205 | -0.577 | **0.998** |
| CcHyogo - CcKyoto == 0 | 1.999 | 2.325 | 0.860 | 0.982 |
| CcShimane - CcKyoto == 0 | 2.105 | 2.175 | 0.968 | 0.966 |
| CcfcTokyo - CcKyoto == 0 | -11.322 | 2.325 | -4.871 | **< 0.001** |
| CcfcKyoto - CcKyoto == 0 | -7.426 | 0.619 | -11.996 | **< 0.001** |
| CcfcHyogo - CcKyoto == 0 | -9.541 | 2.302 | -4.144 | **< 0.001** |
| CcfcShimane - CcKyoto == 0 | -5.991 | 2.166 | -2.765 | 0.071* |
| CcShimane - CcHyogo == 0 | 0.106 | 2.172 | 0.049 | 1.000 |
| CcfcTokyo - CcHyogo == 0 | -13.321 | 2.433 | -5.475 | **< 0.001** |
| CcfcKyoto - CcHyogo == 0 | -9.425 | 2.345 | -4.019 | **<0.01** |
| CcfcHyogo - CcHyogo == 0 | -11.540 | 0.631 | -18.294 | **< 0.001** |
| CcfcShimane - CcHyogo == 0 | -7.990 | 2.185 | -3.657 | **<0.01** |
| CcfcTokyo - CcShimane == 0 | -13.427 | 2.255 | -5.954 | **< 0.001** |
| CcfcKyoto - CcShimane == 0 | -9.530 | 2.187 | -4.357 | **< 0.001** |
| CcfcHyogo - CcShimane == 0 | -11.645 | 2.163 | -5.383 | **< 0.001** |
| CcfcShimane - CcShimane == 0 | -8.096 | 0.745 | -10.864 | **< 0.001** |
| CcfcKyoto - CcfcTokyo == 0 | 3.896 | 2.309 | 1.688 | 0.607 |
| CcfcHyogo - CcfcTokyo == 0 | 1.781 | 2.384 | 0.747 | 0.992 |
| CcfcShimane - CcfcTokyo == 0 | 5.331 | 2.226 | 2.394 | 0.178 |
| CcfcHyogo - CcfcKyoto == 0 | -2.115 | 2.316 | -0.913 | 0.975 |
| CcfcShimane - CcfcKyoto == 0 | 1.435 | 2.174 | 0.660 | 0.996 |
| CcfcShimane - CcfcHyogo == 0 | 3.550 | 2.168 | 1.637 | 0.643 |

Table S10. Multiple Comparisons of Means in stomata density: Tukey Contrasts. Boldface indicates significant effects ($P < 0.001$, $P < 0.01$, and $P < 0.05$).

| Linear Hypotheses | Estimated coefficient | Standard error | *z* value | *P* |
|---|---|---|---|---|
| CcKyoto - CcTokyo == 0 | -17.767 | 6.661 | -2.667 | 0.119 |
| CcHyogo - CcTokyo == 0 | -22.520 | 7.980 | -2.822 | 0.080 |
| CcShimane - CcTokyo == 0 | -19.537 | 7.142 | -2.735 | 0.100 |
| CcfcTokyo - CcTokyo == 0 | 9.238 | 4.407 | 2.096 | 0.388 |
| CcfcKyoto - CcTokyo == 0 | -7.755 | 6.483 | -1.196 | 0.922 |
| CcfcHyogo - CcTokyo == 0 | 5.077 | 7.387 | 0.687 | 0.997 |
| CcfcShimane - CcTokyo == 0 | 2.018 | 6.877 | 0.293 | 1.000 |
| CcHyogo - CcKyoto == 0 | -4.753 | 6.909 | -0.688 | 0.997 |
| CcShimane - CcKyoto == 0 | -1.770 | 6.395 | -0.277 | 1.000 |
| CcfcTokyo - CcKyoto == 0 | 27.005 | 6.898 | 3.915 | **<0.01** |
| CcfcKyoto - CcKyoto == 0 | 10.011 | 4.407 | 2.272 | 0.284 |
| CcfcHyogo - CcKyoto == 0 | 22.844 | 6.558 | 3.484 | **<0.05** |
| CcfcShimane - CcKyoto == 0 | 19.784 | 6.281 | 3.150 | **<0.05** |
| CcShimane - CcHyogo == 0 | 2.983 | 6.399 | 0.466 | 1.000 |
| CcfcTokyo - CcHyogo == 0 | 31.758 | 8.401 | 3.780 | **<0.01** |
| CcfcKyoto - CcHyogo == 0 | 14.765 | 7.203 | 2.050 | 0.418 |
| CcfcHyogo - CcHyogo == 0 | 27.597 | 4.481 | 6.159 | **< 0.001** |
| CcfcShimane - CcHyogo == 0 | 24.538 | 6.522 | 3.762 | **<0.01** |
| CcfcTokyo - CcShimane == 0 | 28.775 | 7.487 | 3.843 | **<0.01** |
| CcfcKyoto - CcShimane == 0 | 11.782 | 6.572 | 1.793 | 0.594 |
| CcfcHyogo - CcShimane == 0 | 24.614 | 6.255 | 3.935 | **<0.01** |
| CcfcShimane - CcShimane == 0 | 21.555 | 4.843 | 4.451 | **< 0.001** |
| CcfcKyoto - CcfcTokyo == 0 | -16.993 | 6.659 | -2.552 | 0.157 |
| CcfcHyogo - CcfcTokyo == 0 | -4.161 | 7.751 | -0.537 | 0.999 |
| CcfcShimane - CcfcTokyo == 0 | -7.220 | 7.184 | -1.005 | 0.969 |
| CcfcHyogo - CcfcKyoto == 0 | 12.832 | 6.764 | 1.897 | 0.522 |
| CcfcShimane - CcfcKyoto == 0 | 9.773 | 6.404 | 1.526 | 0.770 |
| CcfcShimane - CcfcHyogo == 0 | -3.059 | 6.290 | -0.486 | 1.000 |

Table S11. Generalized linear mixed model (GLMM) dataset to show the relationship between the ratio of developed land area and stomata size or density. Boldface indicates significant effects (P < 0.001, P < 0.01, and P < 0.05).

| Response variable / Explanatory variable | Estimated coefficient | Standard error | *z* value | *P* |
|---|---|---|---|---|
| Stomata size/ | | | | |
| (Intercept) | 48.297 | 1.531 | 31.556 | **<0.001** |
| The ratio of developed land area | -0.019 | 0.018 | -1.066 | 0.287 |
| Kyoto | 3.975 | 1.916 | 2.075 | **<0.05** |
| Hyogo | 4.979 | 1.933 | 2.576 | **<0.01** |
| Shimane | 5.298 | 1.911 | 2.772 | **<0.01** |
| Stomata size/ | | | | |
| (Intercept) | 52.224 | 3.216 | 16.240 | **<0.001** |
| The ratio of developed land area | -0.028 | 0.038 | -0.738 | 0.461 |
| Kyoto | -7.932 | 4.025 | -1.971 | **<0.05** |
| Hyogo | -12.461 | 4.062 | -3.068 | **<0.01** |
| Shimane | -10.733 | 4.016 | -2.673 | **<0.01** |

Table S12. Generalized linear mixed model (GLMM) dataset to show the effects of leaf area and sampling date in Cc. Stomata size was the response variable, region identity, leaf area, and sampling date were the explanatory variables, and study site identity was random term. Boldface indicates significant effects (P < 0.01 and P < 0.05).

| Response variable / Explanatory variable | Estimated coefficient | Standard error | *z* value | *P* |
|---|---|---|---|---|
| Stomata size/ | | | | |
| (Intercept) | 230.367 | 80.591 | 2.858 | **<0.01** |
| Leaf area | 0.362 | 1.832 | 0.198 | 0.843 |
| Kyoto | -17.168 | 13.126 | -1.308 | 0.191 |
| Hyogo | -28.966 | 16.011 | -1.809 | 0.070 |
| Shimane | -13.139 | 8.710 | -1.508 | 0.131 |
| Time (sampling date) | -0.922 | 0.405 | -2.279 | **<0.05** |
| Leaf area and Kyoto | -3.463 | 2.442 | -1.418 | 0.156 |
| Leaf area and Hyogo | 0.219 | 1.937 | 0.113 | 0.910 |
| Leaf area and Shimane | 0.754 | 2.341 | 0.322 | 0.747 |

Table S13. Comparison of pollen size (pollen area) in Cc and Ccfc. Pollen area was measured by Fiji (Schindelin *et al*., 2012). 100 pollens were collected and counted from one stamen in Cc and Ccfc, respectively. Different letters behind the value indicate a significant difference ($p < 0.05$).

| Species | Pollen area (Average ± SE) $\mu m^2$ |
|---|---|
| *C. communis* | 3892.65 ± 42.83 A |
| *C. c.* f. *ciliata* | 2766.74 ± 28.97 B |

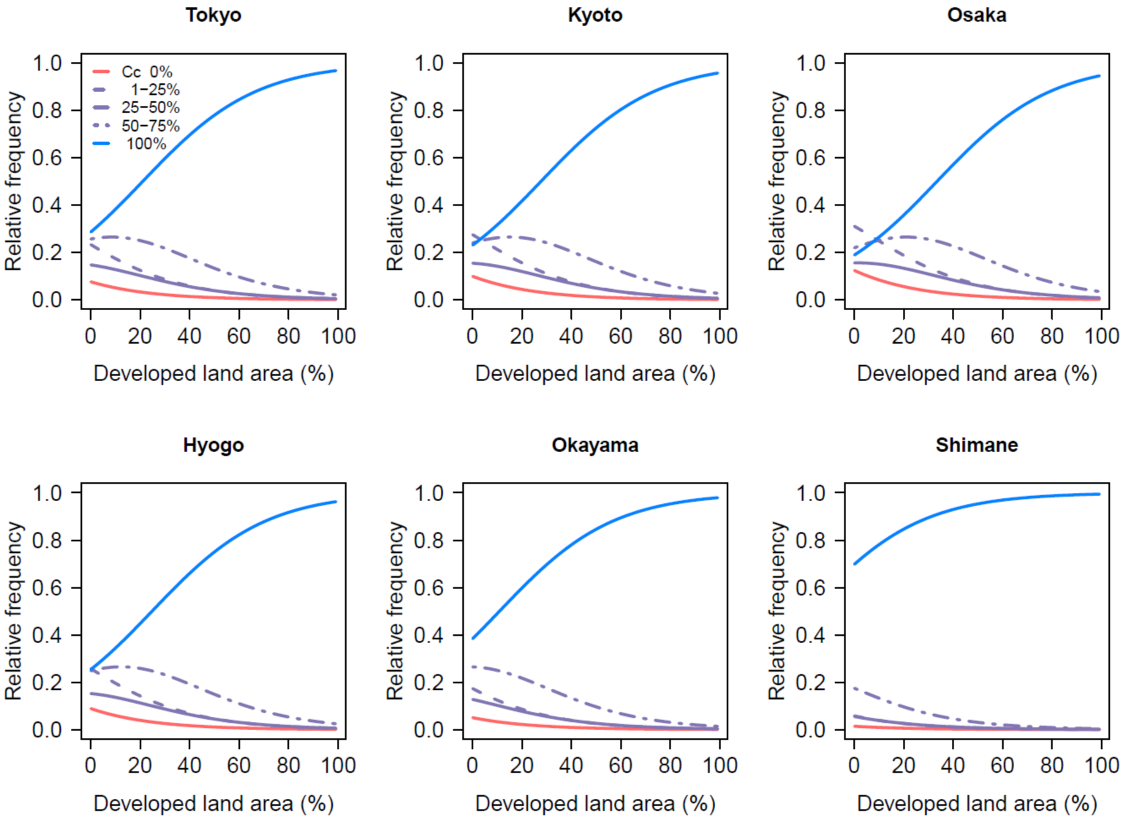


**Fig. S1.** Relationship between the developed land area and relative frequency of the distribution ratio. Legend color is identical to the distribution ratio of *Commelina communis* (Cc) (red, purple, and blue: 1, 2-4, 5); the same meaning as 1: 100% of *C. c.* f. *ciliata* (Ccfc), 2: 50-75%, 3: 25-50%, 4: 1-25%, 5: 0%. Relative frequency indicates the prediction of plant distribution patterns. This model is based on the data of ordinal logistic regression analysis and the model with spatial scale 1000 m was selected due to the lowest AIC value (Table S6). The horizontal axis indicates the percentage of developed land areas in each site. The vertical axis indicates the relative frequency of the distribution pattern in each developed land area rate. For example, when the ratio of developed lands is 20%, the relative frequency of only Cc growing (distribution ratio: 5) seems to be around 60%.

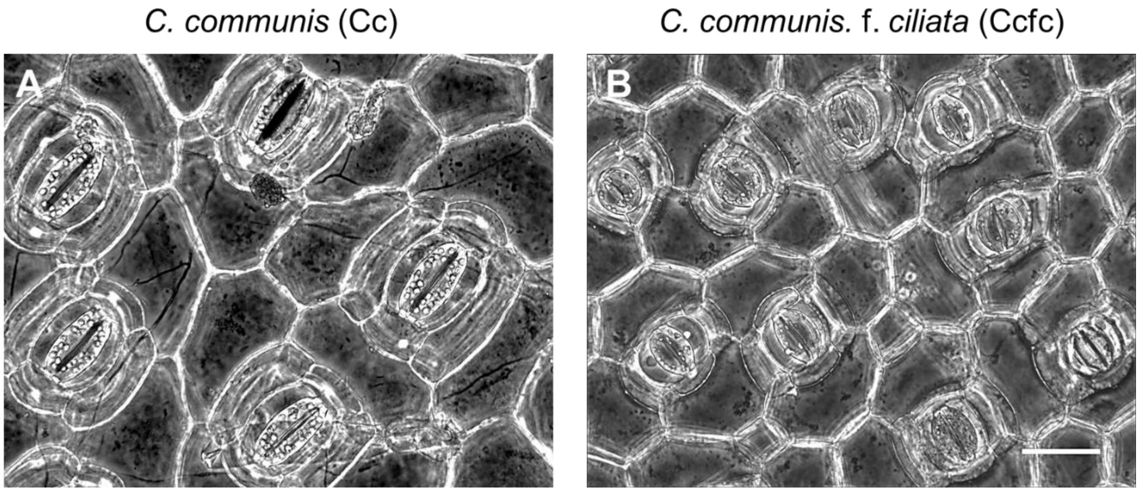


**Fig. S2**. Comparison of stomata in Cc and Ccfc. Stomata of *Commelina communis* (Cc) and *C. c.* f. *ciliata* (Ccfc) were shown in A and B, respectively. Both samples were collected from rural area in Hyogo. Scale bar: 50 µm.

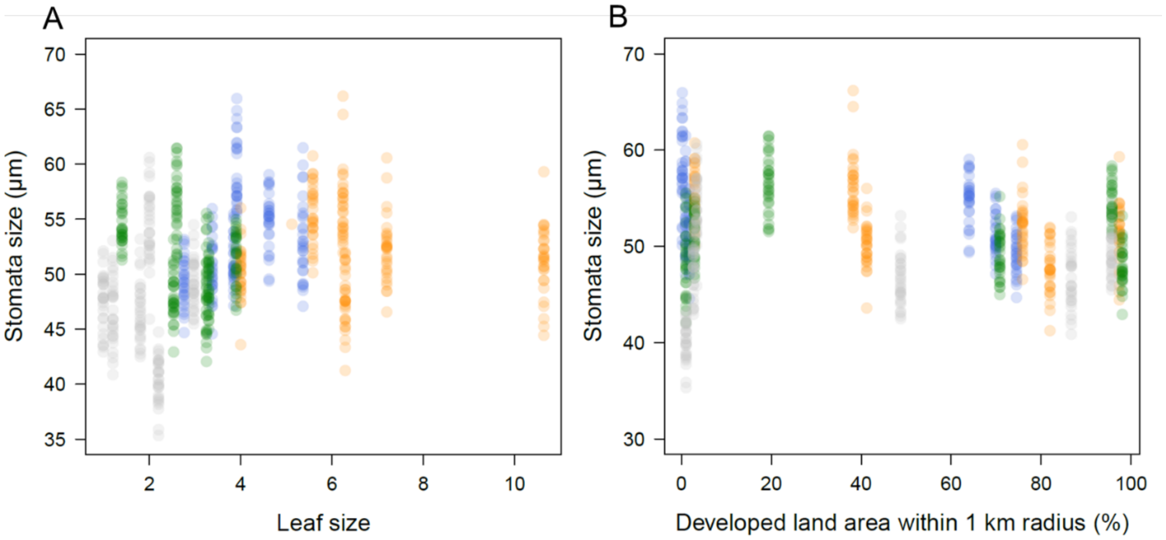


**Fig. S3.** The relationship between stomata size and the developed land area. Each color indicates the region; silver, green, orange, and blue are Tokyo, Kyoto, Hyogo, and Shimane, respectively. (A) Leaf area and stomata size to show the existence of region units. Leaf sizes were calculated by multiplying leaf width and leaf length. (B) The ratio of developed land area and stomata size. Figure S3A showed that there were some tendencies of stomata size depending on regions, which indicated that the comparison of regions was meaningful.

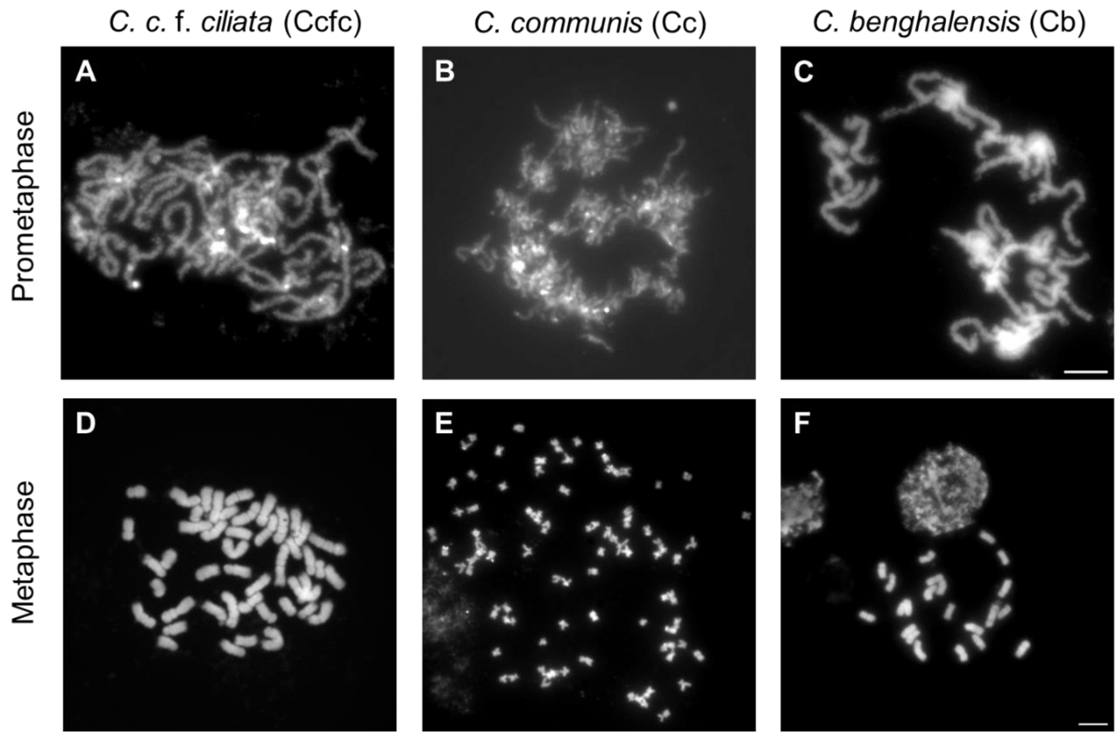


**Fig. S4.** Mitotic prometaphase and metaphase of Cc, Ccfc, and Cb. (A) Prometaphase of Ccfc chromosomes stained by DAPI. (B) Prometaphase of Cc chromosomes stained by DAPI. (C) Prometaphase of Cb chromosomes stained by DAPI. (D) Metaphase of Ccfc chromosomes stained by DAPI. (E) Metaphase of Cc chromosomes stained by DAPI. (F) Metaphase of Cb chromosomes stained by DAPI. Scale bar: 5 μm.

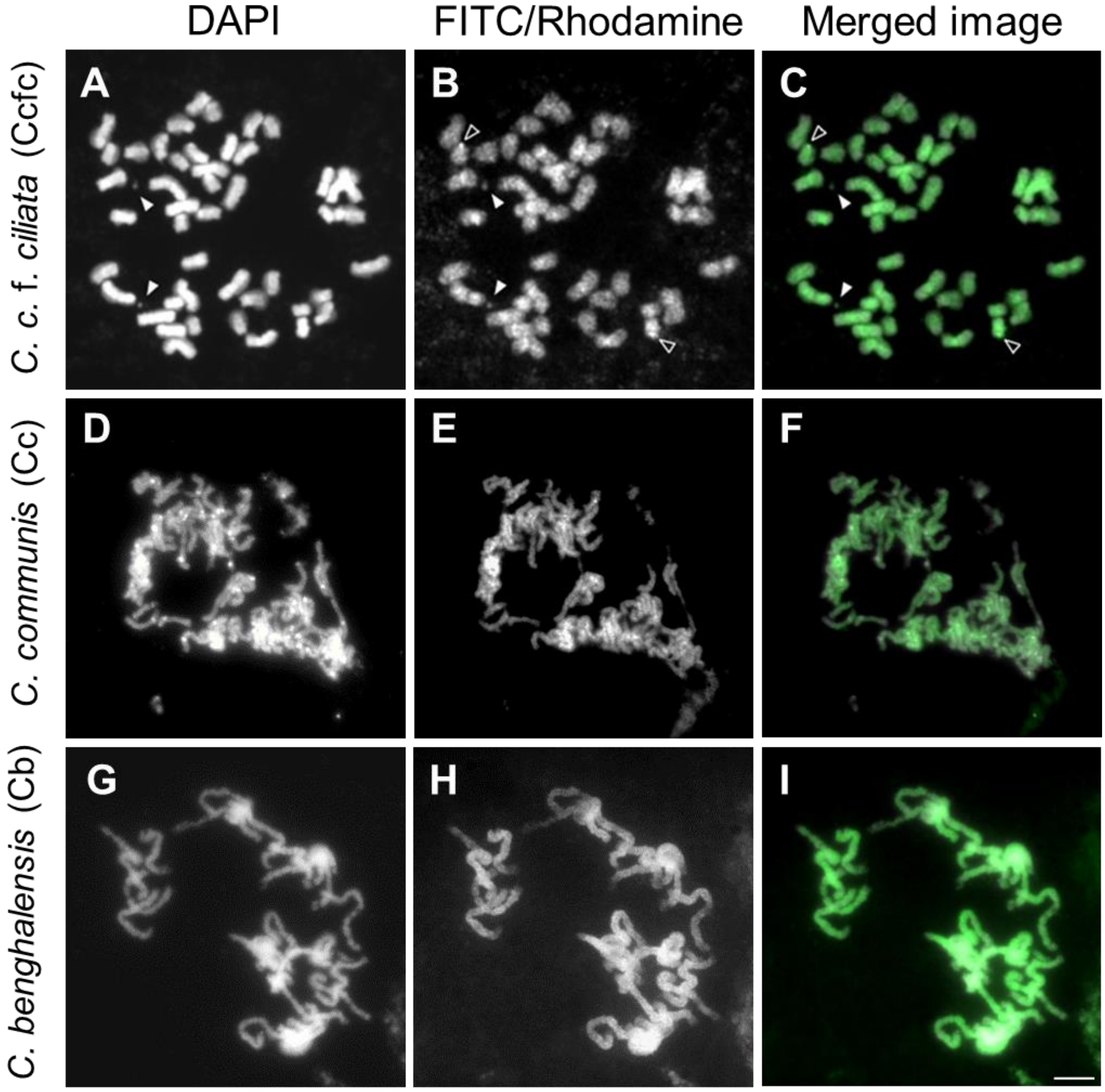


**Fig. S5.** Mitotic metaphases of Cc, Ccfc, and Cb after genomic fluorescence *in situ* hybridization (GISH) using genomic Cb DNA (green). (A) Ccfc chromosomes stained by DAPI. (B) Cb DNA signals labeled by FITC. (C) Cb DNA signals mapped on Ccfc chromosomes (green). (D) Cc chromosomes stained by DAPI. (E) Cb DNA signals labeled by FITC. (F) Cb DNA signals mapped on Cc chromosomes (green). (G) Cb chromosomes stained by DAPI. (H) Cb DNA signals labeled by FITC. (I) Cb DNA signals mapped on Cb chromosomes (green). Arrowheads (white) demonstrate satellite chromosomes. Arrowheads (black) indicate the strong signal patterns. Scale bar: 5 μm.
